# Integrating predictors of host condition into spatiotemporal multi-scale models of virus shedding

**DOI:** 10.1101/2024.10.24.616488

**Authors:** Andrew M Kramer, Christina L Faust, Adrian A. Castellanos, Ilya R Fischhoff, Alison J. Peel, Peggy Eby, Manuel Ruiz-Aravena, Benny Borremans, Raina K. Plowright, Barbara A Han

## Abstract

Understanding where and when pathogens occur in the environment has implications for reservoir population health and infection risk. In reservoir hosts, infection status and pathogen shedding are affected by processes interacting across different scales: from landscape features affecting host location and transmission to within-host processes affecting host immunity and infectiousness. While uncommonly done, simultaneously incorporating processes across multiple scales may improve pathogen shedding predictions. In Australia, the black flying fox (*Pteropus alecto*) is a natural host for the zoonotic Hendra virus, which is hypothesized to cause latent infections in bats. Re-activation and virus shedding may be triggered by poor host condition, leading to virus excretion through urine. Here, we developed a statistical modeling approach that combined data at multiple spatial and temporal scales to capture ecological and biological processes potentially affecting virus shedding. We parameterized these models using existing datasets and compared model performance to under-roost virus shedding data from 2011-2014 in 23 roosts across a 1200-km transect. Our approach enabled comparisons among multiple model structures to determine which variables at which scales are most influential for accurate predictions of virus shedding in space and time. We identified environmental predictors and temporal lags of these features that were important for determining where reservoirs are located and multiple independent proxies for reservoir condition. The best-performing multi-scale model delineated periods of low and high virus prevalence, reflecting observed shedding patterns from pooled under-roost samples. Incorporating regional indicators of food scarcity enhanced model accuracy while incorporating other stress indicators at local scales confounded this signal. This multiscale modeling approach enabled the combination of processes from different ecological scales and identified environmental variables influencing Hendra virus shedding, highlighting how integrating data across scales may improve risk forecasts for other pathogen systems.

## INTRODUCTION

The presence and abundance of pathogens in a location result from physical, ecological, and physiological processes occurring across multiple biological scales. Pathogens infect cells within individual hosts; pathogens are transmitted between individuals; and the geographic distribution of a host population dynamically responds to changes in regional climate and local environmental conditions, which combine to influence resource availability and feedback to affect host condition and distribution. Multi-scale models have helped identify biological scales contributing disproportionately to transmission dynamics (Orton et al. 2020, Tsao et al. 2020) and informed disease intervention and control (Guo et al. 2015). While interacting scales underpin infection dynamics observed in natural systems, there are no standard methods for incorporating multiscale processes in disease ecology. Researchers have implemented multi-scale models using various model types (dynamical, statistical) and linkages (correlational, mechanistic) between scales (Hasenauer et al. 2015, Childs et al. 2019, Kramer et al. 2019). Despite the potential for multi-scale models to offer new insights, linking spatially and temporally dynamic processes across biological scales remains challenging partly due to the data necessary to characterize feedbacks and dependencies in complex systems (Garabed et al. 2019).

An illustrative example of this complexity is the interaction between individual body condition and infection, which can, in turn, impact epidemic dynamics across a population (Beldomenico et al. 2008, Beldomenico and Begon 2010). While there are several condition metrics (Jakob et al. 1996, Peig and Green 2009), body condition generally describes the energetic status of an individual with direct implications for fitness (survival & reproduction). While host condition responds to processes happening at several scales, most studies investigating links between body condition and infection status find negative associations (Sanchez et al. 2018). Animals in poor body condition (e.g., from low resource input) may divert energy from immune responses needed to prevent or control infections (Plowright et al. 2024). For example, poor nutrition is associated with an increased amount and duration of virus shedding in a songbird species (Owen et al. 2021). Infections can also reduce host condition through anorexia or investment in immunity (Kyriazakis et al. 1998, Reeder et al. 2012, Verant et al. 2014).

Many wildlife diseases lack sufficient temporal and spatial sampling to develop multi-scale models. However, decades of Hendra virus research make it a valuable system for building multi-scale models and testing key hypotheses. Hendra virus (HeV) is a Henipavirus that circulates in flying fox populations in eastern Australia, and the virus is shed and transmitted through urine. Because the virus can spill over to domesticated horses and humans (Murray et al. 1995, Rogers et al. 1996), extensive research efforts have been carried out to identify drivers of HeV in wild flying fox populations (Field et al. 2001). Black flying foxes (*Pteropus alecto*) are a primary reservoir for HeV (HeV-g1) and are likely source of the majority of spillover cases (Edson et al. 2015, Annand et al. 2022, Peel et al. 2022). Research on HeV shows complex transmission patterns with significant variation in the prevalence and timing of HeV shedding (Plowright et al. 2015). Once shed, HeV is not viable in the environment for more than a day (Fogarty et al. 2008, Martin et al. 2015), so identifying where reservoir hosts are on the landscape is essential for understanding virus shedding patterns. The occurrence and abundance of black flying foxes have previously been linked to differences in virus shedding (Paez et al. 2017); however the high mobility of flying foxes makes it challenging to match viral prevalence and population data in space and time across their range. Flying foxes exhibit nomadic behavior to track dynamic flowering of native Eucalyptus and other Myrtaceae species, which involves frequent switching of roosts hundreds of kilometers apart within a month (Palmer et al. 2000, Welbergen et al. 2020). HeV shedding is often measured from pooled under-roost urine samples and is summarized as prevalence (percentage of samples positive). HeV prevalence at a roost ranges from undetectable levels to over 60% (Edson et al. 2015, Field et al. 2015, Peel et al. 2019). A leading hypothesis is that environmental stress increases HeV shedding, driving observed spillovers (Plowright et al. 2015).

Even though host condition is recognised to impact population-level processes, most studies use individual-level data, such as fat scores and stress biomarkers, to link host condition to infection status or severity. Instead, using population-level indicators of resource availability may highlight environmental conditions that periodically enhance host susceptibility, reactivate latent infections, or increase pathogen shedding. Directly measuring physiological stress and body condition in wild flying foxes is extremely difficult. Here, we tested whether models of population-level stress and condition, based on three distinct proxies for physiological stress, can approximate the spatiotemporal variation in host condition that influences virus shedding. For example, rehabilitation intake numbers are likely to reflect changes in energetics and body condition, with the rate of admissions to rehabilitation facilities across eastern Australia serving as a proxy of population-level stress (Mo et al. 2020). Previously, nutritional stress caused by acute food shortages in winter and spring has been linked to preceding El Niño events (Becker et al. 2023, Eby et al. 2023), and unfavorable weather conditions in the preceding year have been correlated with higher shedding pulses (Paez et al. 2017). Additionally, in response to acute food shortages, flying foxes form new fissioned roosts outside their historical range (Eby et al. 2023).

Using HeV as a case study, we developed a multiscale modeling approach that incorporated factors hypothesized to influence both the dynamic distribution and body condition of reservoir hosts in space and time. These models took a statistical mining approach to combine empirical relationships from separate data sources that did not contain information about HeV infection but could act as proxy indicators of biological processes such as food availability, roost selection, and physical condition. The model was designed to estimate spatiotemporally dynamic risk of virus prevalence based on distinct statistical signals that may describe variation in virus shedding. In contrast to approaches that combine all environmental predictors into a single model (implicitly spanning all scales simultaneously), our approach enhanced interpretability by linking specific proxies to *a priori* hypotheses about the mechanisms driving virus shedding. This process could be described as hypothesis-driven feature construction. We validated model predictions using historical virus prevalence data from black flying fox roosts. The modular nature of our new multiscale modeling approach supports the integration of new data and new models as system knowledge expands.

## METHODS

### General approach

We developed four component models using independent data sets of observed locations and potential indicators of physiological stress among reservoir hosts (full details in Supporting Information). These components occurred at different spatial scales and were combined and scaled to provide a spatially and temporally explicit prediction of virus shedding (summarised as prevalence) that could be compared to field data.

### Case Study

Hendra virus component models. We predicted the risk of HeV shedding in subtropical eastern Australia by linking component models split across two scales to 1) identify where flying fox reservoir species are likely to occur and 2) predict stress proxies for flying fox condition at those locations (Fig. 1). Importantly, we tested whether host condition measured by proxies affected the likelihood of HeV shedding (Fig. 1). We limited our analysis to a region of eastern Australia encompassing all confirmed HeV spillover events and all roost locations used to train the model on black flying fox presence. This area included the distribution of *Pteropus alecto* in coastal subtropical and tropical regions (Hall and Richards 2000, Churchill 2009).

**Figure 1.**
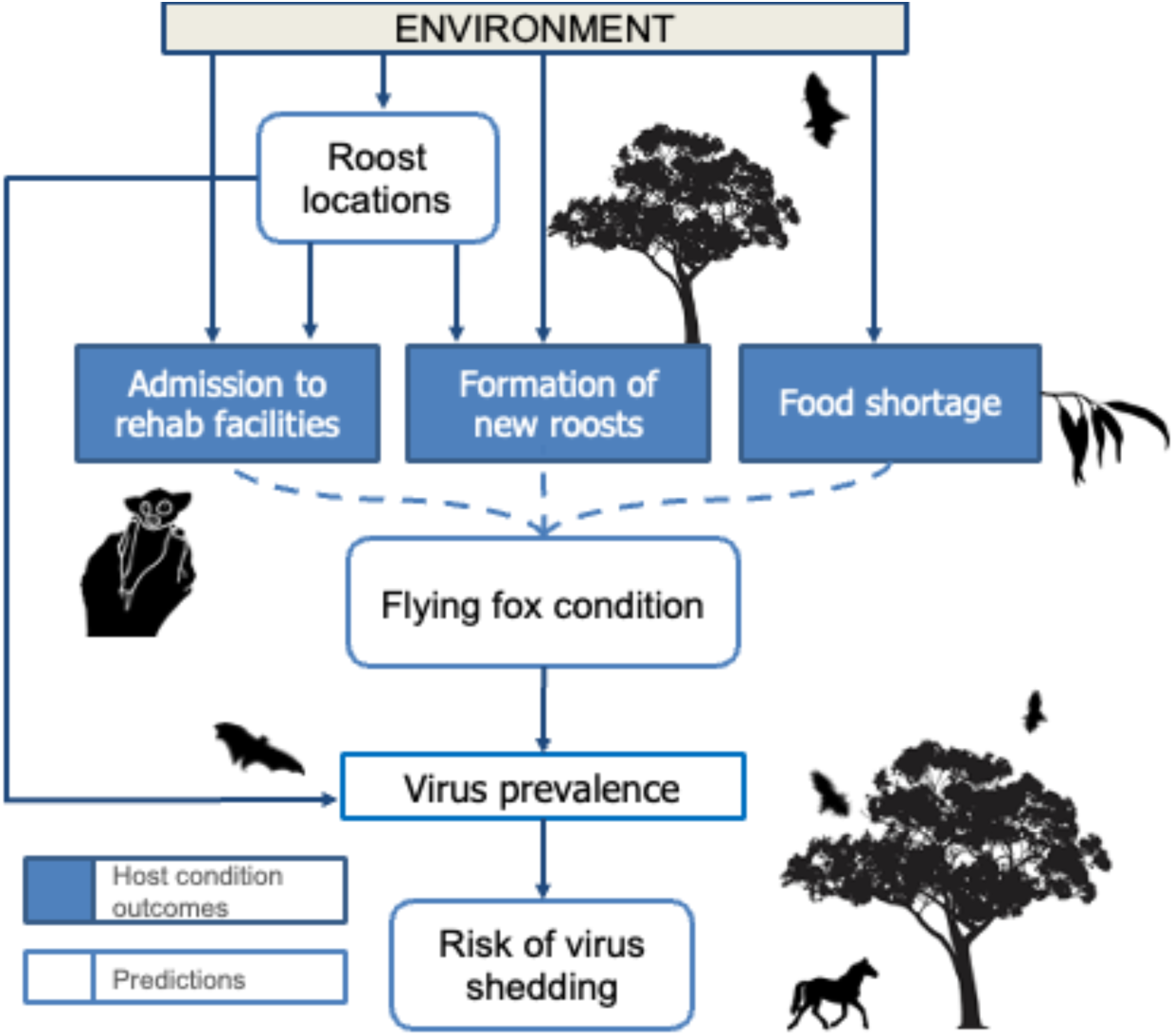
Conceptual multiscale model of Hendra virus shedding. Detecting virus shedding in individual bats is challenging due to high costs and logistical difficulties of field studies. However, shedding can be partially estimated by calculating virus prevalence from pooled under-roost urine samples. Virus shedding is hypothesized to increase during, or following, periods when reservoir hosts (black flying foxes) are in poor condition. Host condition is also difficult to measure directly but can be approximated in multiple ways: through the rate of black flying fox admissions to rehabilitation facilities, by the formation and persistence of new roosts on the landscape (this is thought to be initially an acute response to lack of food), and the occurrence of occasional regional food shortages. These three proxies for host condition are influenced by environmental conditions, which also play a role in where bat roosts are located on the landscape. Our multiscale model links statistical predictions of key components (roost locations and flying fox condition) to estimate virus prevalence across space and time, enhancing our understanding of where and when bats are most likely to actively shed virus.

For each component model, we applied generalized boosted regressions to spatiotemporally-explicit environmental covariates reflecting contemporaneous and historical conditions by incorporating 2 to 24-month lags. We selected these lags as many native black flying fox food resources (Palmer et al. 2000, Markus and Hall 2004) do not flower annually and instead produce nectar and pollen in episodic events that reflect lagged environmental conditions (Eby and Law 2008, Hawkins et al. 2018). Flowering can be highly dynamic in certain native species and is thought to be driven in part by cumulative climatic conditions (Law et al. 2000, Birtchnell and Gibson 2006, Hudson et al. 2010). We included the Oceanic Niño Index (ONI), Southern Oscillation Index (SOI), and Southern Annular Mode (SAM) as reliable climatic indicators. Local environmental conditions included summary metrics of temperature, precipitation, and primary productivity on a ≈ 5 km grid. For lagged conditions we used cumulative values or standard deviations to account for variability in conditions. We also incorporated land cover data into models and summarized it as the proportion of land cover within a 20 km radius. In other settings, flying foxes can forage within smaller or larger radii, dependent on available resources (Palmer 1997, Palmer et al. 2000), but 20 km encapsulates a reasonable foraging distance from a roost by individual *P. alecto* (Palmer 1997).

### Reservoir host locations

*Pteropus alecto* is highly social and roosts in colonies. Colony sizes vary significantly throughout the year and among specific roosts (Lunn et al. 2021). We therefore focused on modeling roost occupancy of black flying foxes across the study region using survey data collected in Queensland (2003-2021) and New South Wales (2012-2019) through the National Flying Fox Monitoring Program (NFFMP) (National Flying Fox Monitoring Program, 2020) and grey-headed flying fox population monitoring (Eby et al. 2022a). The NNFMP data provided the most comprehensive dataset (n = 14,952 unique roost observations over 15 years) of flying fox locations in Australia, despite not covering the entire species range and having variation in frequency of counting. We supplemented this data withoverwintering roost locations and observations from 2002-2019 (Eby et al. 2022a), creating a total of 18,861 records. We disaggregated locations to the monthly time scale such that a roost presence was marked for each grid cell when it was known to be occupied by at least one black flying fox (n = 10,622). We used roost observations without black flying foxes (n= 8,239) as absence points. We then fit a boosted regression tree model using the gbm package in R (Greenwell et al. 2022).

### Host condition

We identified environmental predictors of three separate proxies for host condition that we hypothesized to reflect stress in this system: rehabilitation admissions, fissioning of roosts, and acute food shortages. We considered that rehabilitation and new roosts are dependent on spatial (x) and temporal (t) variation, whereas food shortage is only temporal (t) within our study area because it usually occurs at geographic scales greater than bat movements. These metrics do not directly measure virus shedding, but based on previous studies we expected increased stress to correlate with higher virus shedding in this system (Becker et al. 2023, Eby et al. 2023). Additional information on each model and the calculation of corrected AUC is available in the Supporting information.

#### Rehabilitation admissions

We modeled the probability of flying fox rehabilitation, P(A_xt_), using monthly intake records from WIRES Mid North Coast (n = 695) and Northern Rivers (n = 1027) in New South Wales (https://figshare.com/s/ddb5a1584609b20f6596). The rehabilitation centers recorded the species, originating postcode, and intake date. We fit a boosted regression tree model to presence and background points to predict the probability of any flying fox being brought to rehabilitation centers. We defined presence data based on flying fox admissions between 2005 to 2020 (n = 5,075) and extracted environmental characteristics from each postcode polygon for the month and year of intake. We chose background data within the same postcodes and weighted random sampling of dates.

#### New Roost Formation

We constructed a boosted regression tree model to predict the probability of a location supporting a new, fissioned overwintering roost: P(R_xt_). Newly established roosts have higher rates of virus shedding (Becker et al. 2023); therefore, identifying environmental features associated with the formation of these new overwintering roosts may provide a valuable indicator of stress. We used the aforementioned dataset on overwintering roosts to identify new black flying fox roosts (n= 195) formed between 2000 and 2019 (Eby et al. 2022a). We used conditions from August (the last month of winter) in the year a new roost was formed, or the last month of acute food shortage if the roost was formed in a year with food shortage, as presence data. We selected background data weighted by predictions from the roost SDM.

#### Food Shortage

To model the probability of acute food shortages, P(F_t_), we used records of nectar shortage (Eby et al. 2022b). Acute food shortage (hereafter referred to as “food shortage”) is a binary response defined every month based on surveys from apiarists over an approximate area of 4000 km^2^ in New South Wales from January 1998 to March 2020. We assumed food shortages were equivalent across the entire study region and developed a model based on only global climatic features (ONI, SOI, SAM). We applied a gradient boosted model (Chen et al. 2023) to predict the probability of an food shortage using a binary response of months with (n = 22) and without (n=245) food shortages.

### Multiscale model of Hendra shedding

We first used the roost occupancy model to locate a temporally dependent number of roosts across the study area. We determined the number of roosts each month based on predictions of a generalized additive model of occupied roosts based on the NFFMP dataset. We randomly distributed the number of roosts for a given month and weighted the probability of choosing a given grid cell using the roost occupancy model output. We did not allow multiple roosts to occupy the same grid cell, resulting in roost spacing of at least 5 km. We did this to ensure roosts are approximately as distant as the minimum observed distance (2 km). We then assigned each roost a unique foraging area encompassing the space closest to that roost with tessellation followed by truncation of any part of a polygon extending more than 50 km from each location. This divided areas between roosts when they were nearby while preventing unrealistically distant areas from being incorporated into roost condition predictions.

We predicted virus shedding as prevalence at a roost location to compare with observed data from pooled under roost sampling. Host condition (C_xt_) predictions were determined by all combinations of three host condition models: P(A) was the probability of bats needing rehabilitation, P(R) was the probability of the site being a new overwintering roost, and P(F) was the probability of an acute food shortage. To quantify the spatial component of stress, we calculated the geometric mean of rehabilitation and new roost formation. We then multiplied the complement of spatial stress by the complement of food shortage, which was expected to affect the entire area and hypothesized to compound spatial stress. Multiplying ensured a cumulative prediction of stress, i.e., low values of one stress type did not offset high values of the others. We combined the results from the three models as follows:

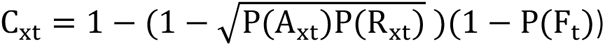

C varies between 1 (when bats are certain to be stressed) and 0 (when bats are unlikely to be stressed). Although this is a naive estimator, we propose it as a reasonable starting point that will be proportional to stress if the component hypotheses are supported. This value was assumed for the entire area within each roost tessellation and represented a naive model for how host condition proxies may interact. For each combination of component models, we considered the maximum possible prevalence to match the highest observed prevalence at a roost in a large-scale Hendra virus survey (MaxP=66.6%; Field et al. 2015). We then calculated the predicted prevalence for each space-time point:

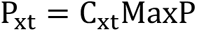

We predicted prevalence across the landscape for a given month by generating 1,000 roost location maps, obtaining the host condition for each distribution, predicting the prevalence within each tessellation, and then averaging those 1,000 random realizations from January 2008 to December 2019. We maximized information gained about how stress proxies vary in space and time by excluding zero prevalences resulting from roost absence from the averages. We incorporated uncertainty in the host condition models by bootstrapping the input data for each model 1,000 times and refitting to the bootstrapped data. Therefore, each roost location map for a month was associated with one of the bootstrapped host condition models.

### Model validation and comparison

We compared prevalence predictions from each multi-scale model to observed HeV prevalence estimates from under-roost sampling of 26 roosts across eastern Australia (Field et al. 2015). We only used roosts sampled at least five times within the study and observed to have *P. alecto* within the study period, resulting in 352 observations of 23 roosts from July 2011 to November 2014. Predicted prevalence values for each observed roost were calculated from model predictions for a 20 km buffer around the site coordinates.

Host condition can be affected by both acute and chronic effects. To determine the appropriate temporal scale for linking host condition to virus shedding, we first considered cumulative lagged conditions by comparing the 3 to 12-month averages of each condition to contemporaneous condition prevalence predictions. For each component model, we selected the most informative lag based on Spearman correlation coefficients of predictions against observations. Using the cumulative condition lags that maximized this correlation coefficient, we investigated the influence of each component model and all possible combinations on prevalence predictions.

We also considered a null model that used only the roost suitability predictions (no condition); the null model tested whether the likelihood of host presence was sufficient to explain variation in prevalence. This resulted in eight sets of predictions, and we compared the relative performance of each of these predictions using root mean squared error (RMSE) and took the mean of this metric for each sampled roost. The lowest values of RMSE corresponded to the best-fitting models. Spearman rank correlations for the same set of models provided a complementary measure of relative agreement between predictions and observations.

## RESULTS

Black flying fox roost occupancy and indicators of flying fox condition - measured as rehabilitation admission, new roost formation, and food shortage - exhibited predictable spatiotemporal variation across eastern Australia. We linked these component models to predict virus shedding and demonstrate empirical relationships between host locations and conditions with implications for HeV shedding.

### Component model performance

#### Reservoir host locations

Environmental features predicted black flying fox roost occupancy (Fig. 2). Our best model of roost locations had a mean corrected AUC of 0.867. Full details of all model evaluation and formulation are in the Supporting information. The most important variables predicting roost occupation included percentage of land cover (including urban, pasture, cropland, and forest), standard deviations of temperature (maximum, minimum, and range), and cumulative solar exposure. Black flying foxes were more likely to occupy roosts in areas with high percentages of urban land cover, intermediate pasture, low crop cover, both low or high percentage forest cover within foraging radii (20 km), low temperature variability within the last nine months, and higher soil moisture.

**Figure 2.**
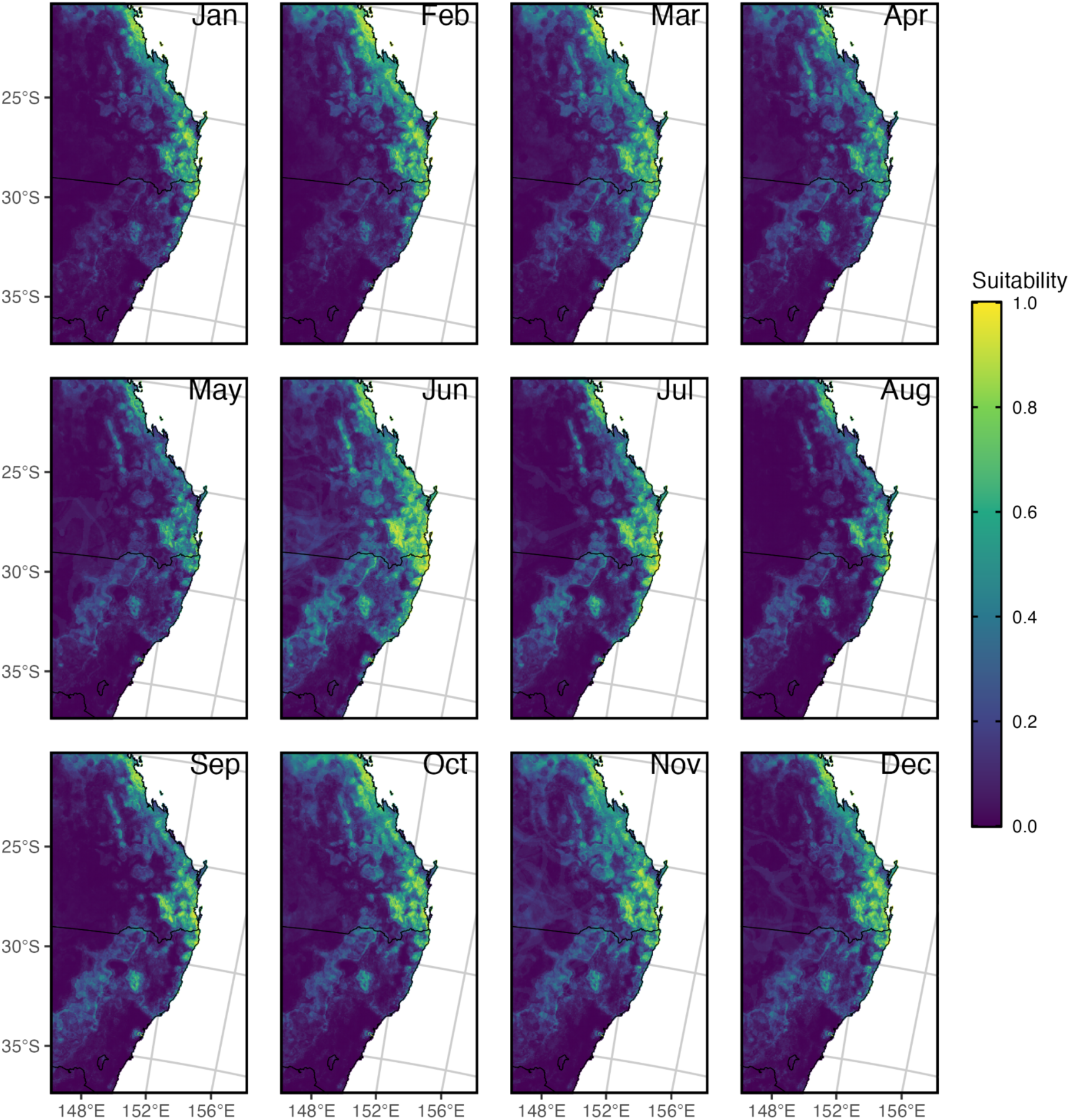
Mean environmental suitability for black flying fox roosts from 1996-2021. Warmer colors show higher predicted suitability to environmental conditions, while cooler colors show lower predicted suitability. Overall, we see consistency in suitable regions for black flying foxes, with the edges of these regions expanding or retracting slightly according to seasonal shifts.

#### Host condition

Full model results for the three proxies of host condition, including partial dependence plots and relative importance scores of the top variables, are reported in the Supporting Information. Our best model of rehabilitation admissions had a mean corrected AUC of 0.876. Land cover classes were most important in these models followed by various weather and climatic variables. Our best model of new roost formation had a mean corrected AUC of 0.9. The probability of a new roost formation was highly spatially dynamic and depended on land cover features, soil moisture, and various lagged temperature and precipitation variables. Our best model of food shortage had a mean corrected AUC of 0.811 (Fig. 3). The most important predictors were higher ONI 9 months prior, lower ONI 21 months prior, and higher SAM 9 months prior. In contrast to the new roost model, the probability of rehabilitation and food shortage were most dependent on temporally varying features, resulting in large variation in predicted host conditions between months and years.

**Figure 3.**
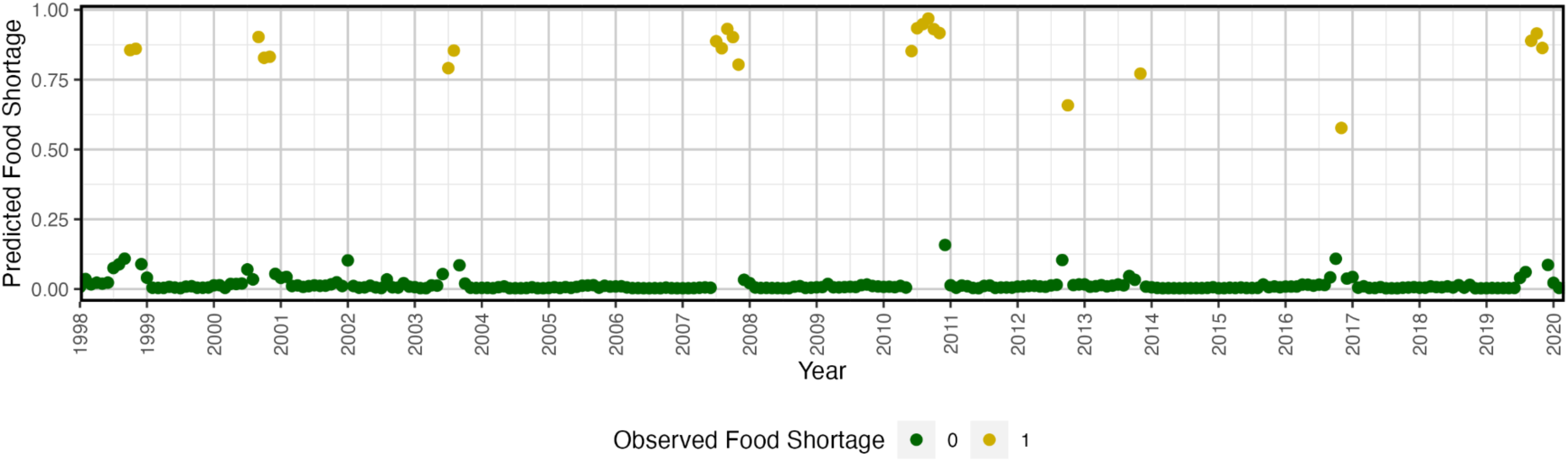
Monthly predicted probabilities of regional food shortage. Months with the highest predicted probabilities of an acute food shortage corresponded to months with observed acute food shortages (yellow).

### Multi-scale model performance in predicting HeV shedding

To assess multi-scale model performance we initially compared different lags of host condition indicators with the empirical estimates of virus shedding. We found that a cumulative 12-month lag in predicted probability of a food shortage (i.e. the average predicted food shortage over the last 12 months) most closely matched the timing and amplitude of HeV shedding observations from 2011-2014. After comparison with other lengths, this cumulative 12-month lag performed the best for our other proxies of stress and was used for all multi-scale model comparisons.

The most accurate multi-scale model included the cumulative 12-month food shortage and roost location models (Fig. 4, Table 1). The null model performed better than models that included the rehabilitation and new roost models without food shortage and the version with all three stress models. Predictions of virus prevalence that included the new roost component model were poor, resulting in negative correlations with observed prevalences (Table 1).

**Figure 4.**
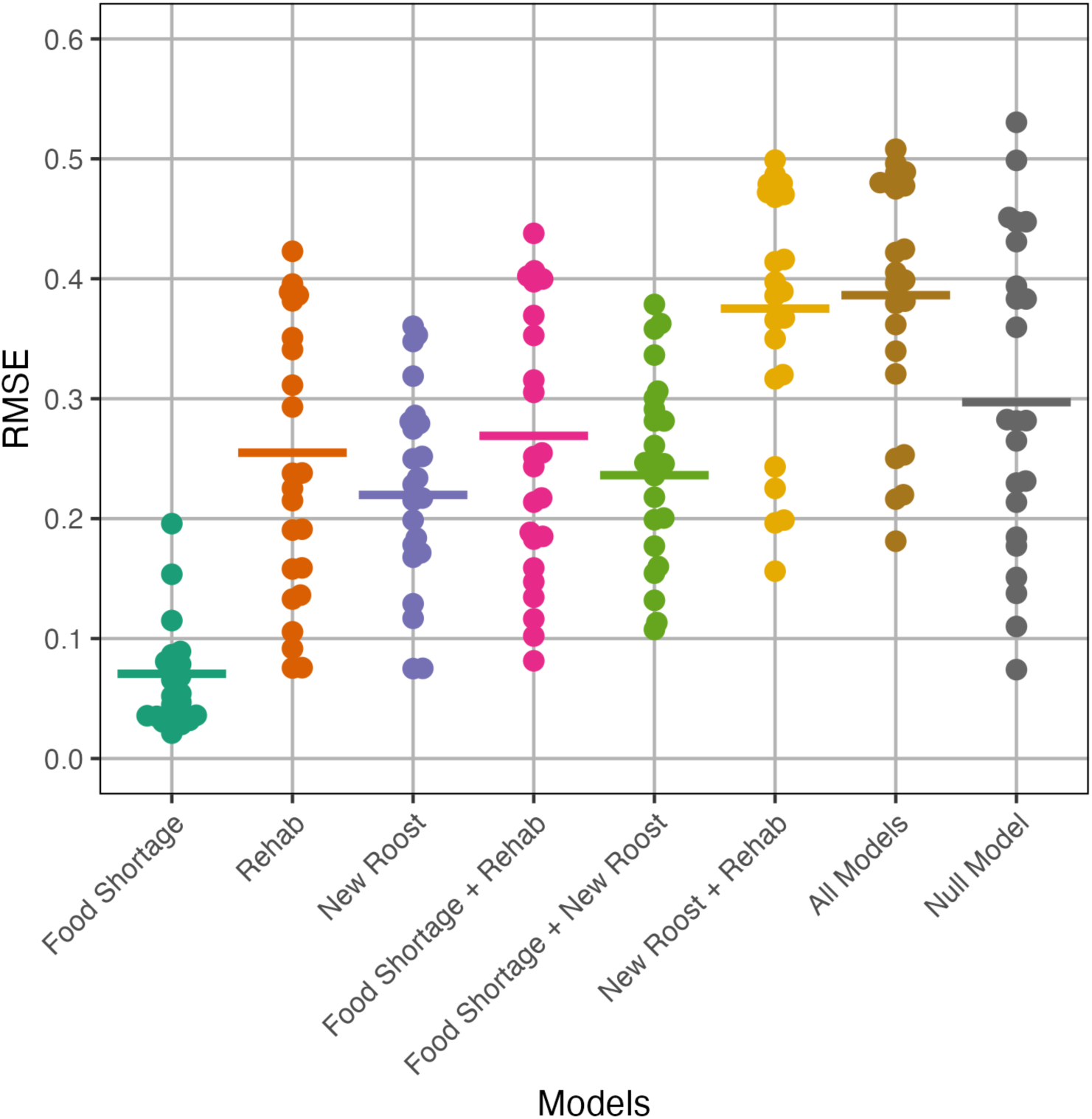
Comparison of predicted to observed prevalence across model structures. Root mean-squared error indicated deviation of predicted values from observations. Lower values of RMSE indicated closer model fits to data. Each point represents the mean RMSE value of a particular roost (N=23 unique roosts). Lines are weighted averages across all roosts accounting for the number of time points observed per roost (ranging from 5 to 45; median 12.5).

**Table 1.** Performance of single and multi-scale model combinations. In the Model column, ‘All condition models’ refers to the multi-scale model with rehabilitation, new roosts, and food shortage models. The null model only includes roost suitability without any condition modifiers. Correlation (to observed shedding prevalence) refers to mean Spearman rank correlation coefficients with the 95% confidence bounds given in parentheses. The values in bold indicate the highest performance.

| Model | RMSE | Correlation |
| --- | --- | --- |
| Food shortage | 0.071 | 0.102<br>(0.097, 0.108) |
| Rehab | 0.255 | 0.002<br>(0.000, 0.010) |
| New Roost | 0.220 | -0.057<br>(-0.059, -0.048) |
| Food shortage + Rehab | 0.269 | 0.042<br>(0.038, 0.048) |
| Food Shortage + New Roost | 0.236 | -0.011<br>(-0.014, -0.003) |
| New Roost + Rehab | 0.375 | -0.042<br>(-0.047, -0.036) |
| All condition models | 0.386 | -0.015<br>(-0.019, -0.009) |
| Null | 0.297 | 0.079<br>(0.072, 0.081) |

When we compared predictions of HeV shedding from 2011-2014, we found the best model predictions differentiated low and high observed prevalence while failing to precisely track the magnitude of observed prevalence (Fig. 5, Supporting information). The best multi-scale model had spatial variation in predictions, but this did not arise from the food shortage component. Instead, spatial variation arose from variation in roost occupancy and tessellations around these roosts in each month. Inclusion of the spatially varying proxies for host condition (rehabilitation intakes and new roost formation) degraded predictions (Table 1). Although the null model (roost occupations only) outperformed some multi-scale models in certain roosts, it was never the best performing (Supporting information).

**Figure 5.**
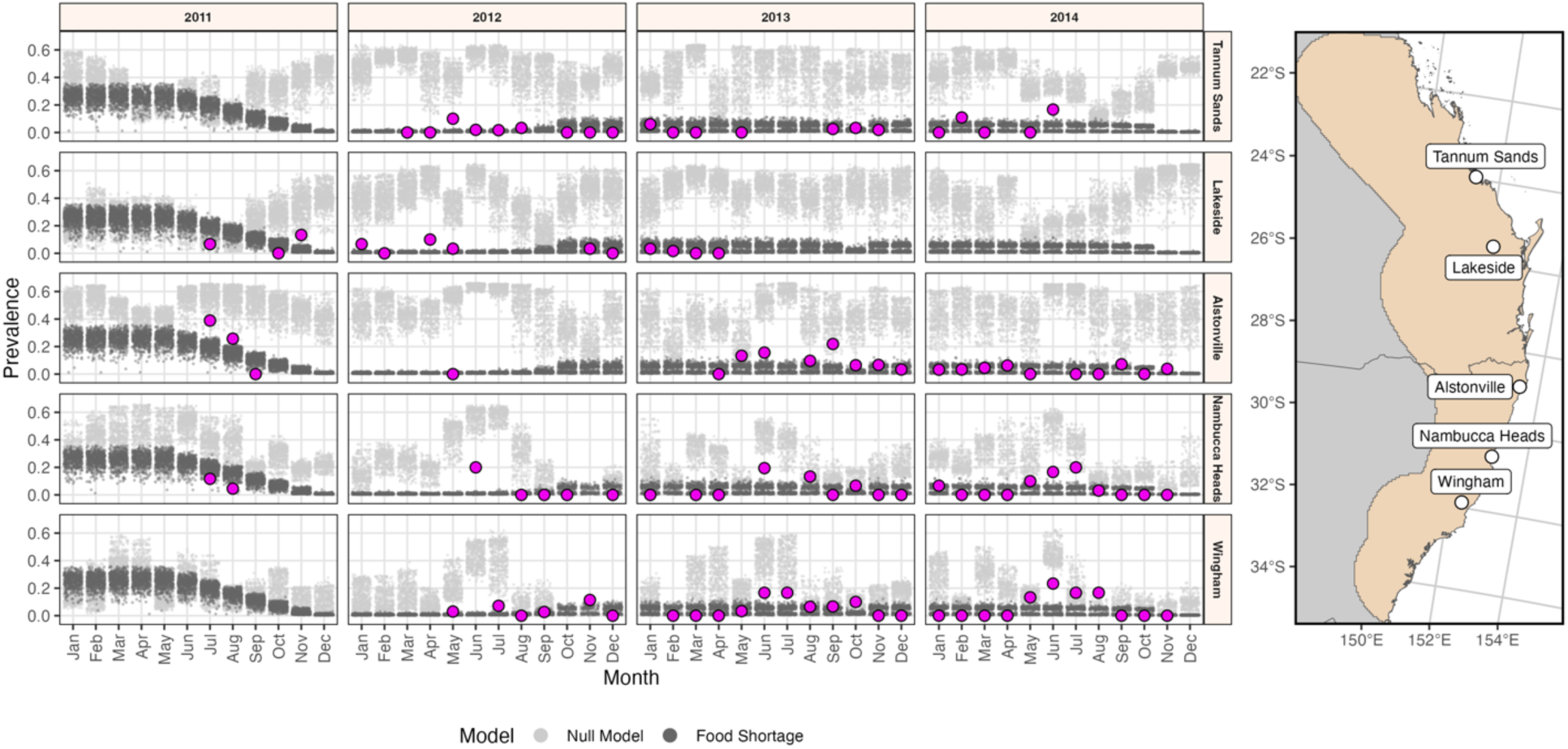
Monthly HeV prevalence for observations compared to null and best-fit model predictions. For a selection of roosts within the study area (beige region of the map), we show the predicted prevalence values for the food shortage model (dark gray points) and the null model based on roost suitability (light gray points). Larger magenta points are observed prevalence values from Field et al. 2015. The panels depict columns as years with observed prevalence values (2011-2014) for sites organized by latitude (rows). Predictions for all sites are in Supporting information.

## DISCUSSION

Our goal was to construct a model that could predict virus shedding by integrating several hypothesized drivers, rather than relying solely on a model closely parameterized to a single dataset. A multi-scale model potentially provides virus shedding predictions based on host location and condition with greater transferability than a universal model directly predicting virus shedding. Even though bats are able to track dynamic resources over long distances, we were able to accurately predict roost suitability across space and time. However, we found that environmental variables explaining location were insufficient to predict virus shedding. Combining the scales encompassing host location and host condition improved spatially and temporally explicit predictions of HeV prevalence and provided support for the hypothesis that stress is linked to virus shedding. Specifically, the superior performance of a model including food shortage provides additional evidence that host condition driven by food shortage is an important driver of virus shedding in this system (Eby et al. 2023).

Although host condition has been proposed as an essential component of transmission, no standard host condition metric exists for inferring impacts on infection outcomes (Sanchez et al. 2018, Vicente-Santos et al. 2023). In this study, we used three proxies for host condition that were agnostic to epidemiological outcomes, available for sufficiently long periods (14-22 years), and from a large spatial area. These types of ecological data - including species occurrences, wildlife rehabilitation admissions, and food availability - could be obtained for other host species and utilized similarly. However, not all three host condition models effectively predict underroost virus prevalence. We found that the host condition model based on the probability of a roost being newly established led to a negative correlation between model predictions and observed data on virus shedding, worse than null model predictions. The poor predictive capability of new roost formation may result from mismatches in time scales from when a new roost forms and when virus shedding increases. Previous studies showed that new overwintering roosts have higher shedding pulses, particularly following an acute food shortage (Becker et al. 2023). The component model we developed only identifies probabilities that a new roost formed, and does not identify ‘new’ overwintering roosts over a longer time period. These new roosts are expected to affect shedding long after they are formed because they are likely in poor foraging locations. The poor fit of this model to observed prevalence data suggests that there is a mismatch in scales of when/where new roosts form and when this is informative for temporal variation of HeV shedding. The new roost host condition model varied with highly spatially variable environmental features (soil moisture, lagged precipitation, lagged temperature), whereas food shortage and rehabilitation models predicted higher temporal variation, and both provided closer predictions to observed data. Nevertheless, the predicted prevalences from the rehabilitation model also exhibited lower correlations with observations than the null model, possibly due to the variety of possible causes for a rehabilitation admission or the limited spatial extent of these data.

Multiscale models including host condition - via food shortages - had superior predictive power, though their predictive performance was modest as measured with correlation (Spearman’s r ≈10%). The better performing models of host condition were not seasonal - this is in contrast to observations of underroost virus shedding, which report seasonal pulses (Paez et al. 2017). Pulses of virus shedding have been linked to seasonal demographic and ecological changes (Wacharapluesadee et al. 2010) - such as birth (Joffrin et al. 2022) and winter food shortages (Becker et al. 2023). We did not observe any seasonality in any of the host condition models, and thus these did not predict seasonal patterns of virus shedding. Furthermore, the lack of seasonality in virus shedding in the multiscale model is also partially driven by the impacts of host condition accumulating over the prior 12 months - which led to predictions of gradual changes in virus shedding. The 12-month time period used was selected based on RMSE in 2011-2014 and, importantly, the fit to observed data was not predicated on any explicit considerations about seasonally dynamic host demography, any data on virus positivity across roosts, or data describing individual-level variation, all of which are known to be influential factors in bat virus shedding (Amman et al. 2012, Dietrich et al. 2018, Peel et al. 2019, Mortlock et al. 2021, Joffrin et al. 2022). Because these processes are not accounted for here, this model can be thought of as improving knowledge of periods of time a pulse is more likely to occur in a location but not the duration of shedding pulses. Future iterations incorporating data on these components and scales may improve predictions and underscore the relative importance of underlying processes contributing to virus shedding.

Model uncertainty may account for limited or absent predictive power for submodels and their combinations. The spatial and temporal span of each dataset differed. There was much more data available for roost locations compared to the limited data available for assessing the signal of bat rehabilitation. These differences are clear in the submodel performances as measured via AUC, and higher uncertainty will necessarily reduce the strength of the correlation between model predictions and prevalence. Future work could incorporate different model structures - e.g., estimating a posterior distribution of condition that can be sampled from to incorporate errors in the multiscale model in a more interpretable manner. Other machine learning tools applied to multiscale biomedical models offer new methods that may be adapted to epidemiological problems (Alber et al. 2019, Peng et al. 2021). Our work and previous studies (i.e. Guo et al. 2015, Kramer et al. 2019, Orton et al. 2020) have highlighted that multiscale models improve predictions of epidemiological outcomes and improve our ability to design more targeted, effective interventions. As new methods and tools are developed for big data, there is an exciting opportunity to apply these to complex systems that operate across multiple ecological scales.

Although HeV presents a uniquely well-studied system for understanding multiscale processes impacting prevalence in a highly mobile host population, there remain limitations to the available datasets. Validation of multiscale models was carried out using a dataset of 23 roosts that had sufficient spatial coverage of the study region but was limited to samples taken over a 41-month period. It is likely that the choice of lags and informative submodels are overfit to this time period and epidemiological patterns. To confirm the multiscale model performs well beyond this period, additional underroost sampling across a large area would be required. Furthermore, the host condition models are based on data that represent a subset of the full geographic range of black flying foxes. Additional data from regions further north in the subtropics and tropics and further south in temperate regions would likely improve predictions of host condition. Finally, if spatial aspects of host condition are more influential than shown here, these could improve prevalence predictions in these regions. This may also allow consideration of whether there is a relevant scale of spatial variation apart from the new roost proxy that performed poorly here.

The modular design of this multiscale approach allows for flexible data integration and consideration of hypothesized drivers of virus shedding acting on different temporal and spatial scales. This contrasts with directly fitting a statistical prevalence model to the many predictors considered across the component models. While a single statistical model might achieve equivalent or superior performance on the validation data, it reduces interpretability, impedes hypothesis generation, and potentially reduces transferability across space and time; aspects for which this approach offers advantages. The statistical integration of multiple models capturing multiple scales can be applied to other systems, as proxies are easily replaced to represent system-specific mechanisms hypothesized to influence pathogen shedding. In this study we present one way for processing, analyzing, and evaluating multi-scale models; this approach may serve as a valuable tool for modeling diseases within their ecological context.

