## Supporting Information for "Integrating predictors of host condition into spatiotemporal multi-scale models of virus shedding"

### Table of Contents:

| | |
|---|---|
| **[A] Overview** | Page 2 |
| **[B] Data description**<br>**[i] Input features**<br>**[ii] Component model labels** | Page 2<br>Page 2<br>Page 6 |
| **[C] Component model details**<br>**[i] Roost distribution**<br>**[ii] Flying fox rehabilitation admissions**<br>**[iii] Formation of new overwintering roosts**<br>**[iv] Acute food shortage** | Page 7<br>Page 8<br>Page 13<br>Page 17<br>Page 21 |
| **[D] Multi-scale model construction & validation**<br>**[i] Roost counts model**<br>**[ii] Model uncertainty**<br>**[iii] Time lags**<br>**[iv] Leave one out description** | Page 24<br>Page 24<br>Page 24<br>Page 24<br>Page 26 |

### [A] OVERVIEW

This supporting information provides more details of the data, analysis and model evaluations to ensure reproducibility of the results presented in the main text. The supporting information is divided into three main sections. The first section describes the data in detail, including: predictor variables used in inputs for component models [B.i] and the data used for constructing each component model and roost count predictions [B.ii]. The second section then presents more information about how component models are constructed along with detailing features that are important in the best fit model for each [C.i-iv]. The last section describes how the multi-scale model is constructed, including roost count predictions [D.i] and uncertainty [D.ii]. The section also includes more details of time lags of component models [D.iii] and comparison of model structures [D.iv].

### [B] DATA DESCRIPTION

#### [B.i] Input features

We collected a variety of environmental variables from publicly available sources. For weather data, we acquired the Australian Gridded Climate Data (AGCD v1), created by the Australian Bureau of Meteorology (BOM). The methodology to create these gridded data are described in detail by (Jones et al. 2009). These data were downloaded at a spatial resolution of around 5 km on a monthly timescale from the years 1996-2021 or as far back in time as they were available (Table S1). Given the importance of previous environmental conditions on eucalypt flowering, we created a series of lagged environmental variables at 2, 3, 6, 9, 12, 15, 18, 21, and 24 months previous. The lags were chosen to match a previous study on nectar production patterns in Australia (Hawkins et al. 2018). For certain variables (precipitation, solar exposure) these lags were calculated as the cumulative value of the variable over these months. Cumulative values provide the cumulative conditions over a particular time period and capture long-term excess or deficit conditions. For others (temperature, precipitation anomaly, NDVI), we calculated the standard deviation of the values over these months. Standard deviation of conditions captures the variance from average conditions and indicates temperatures, vegetation values, etc, that deviate from average conditions that time of year. For example, higher standard deviation of maximum temperatures indicates larger variance of temperatures during that time period.

Previous results showed the importance of a two year lag effect of the Oceanic Niño Index (ONI) on spillover in this study system (Eby et al. 2023). To examine the effects of ONI on our datasets, we collected ONI values for these same lags at each point. Although ONI will not differ spatially for records that occur in the same month and year, these data may inform temporal patterns. We also collected values for the Southern Oscillation Index (SOI) and Southern Annular Mode Index (SAM) to incorporate additional climate indices and match other work investigating nutritional stress in this study system (Lagergren et al. 2023). ONI and SOI values derived from the National Oceanic and Atmospheric Administration's Climate Prediction Center. Values for SAM are derived from methodology described in (Marshall 2003) and come from the British Antarctic Survey.

We collected percentage values of forest, urban areas, crops, and pastures within a 20 km buffer around each record using the terra package (version 1.5-21) in R (Hijmans 2022). These values were calculated using yearly land cover classifications from the Dynamic Land Cover Dataset of Australia (Lymburner et al. 2015). The Dynamic Land Cover Dataset has 22 land cover classes. These classes reflect the structural character of vegetation, and identify cultivated and managed land covers (crops and pastures). We only include four of the 22 land cover types here, so percentages of forest, urban, crops, and pastures do not always add up to 100% for a particular buffer. The choice of a 20 km buffer reflects an approximation of foraging distances from the roost by a *Pteropus alecto* individual (Palmer and Woinarski 1999).

**Table S1. Variables used as environmental features.** Details of component models environmental features were used in (location - roost distribution model, rehab - modeling bats entering wildlife rehabilitation, new - modeling the creation of new roosts, food - modeling periods of food shortage). Data sources include the Australian Gridded Climate Data (AGCD v1), National Oceanic (NOAA), British Antarctic Survey (BAS), and Dynamic Land Cover Dataset (DLCD). Datasets, including derived values, are all deposited in the FigShare (https://figshare.com/s/ddb5a1584609b20f6596).

| Name | Variable Code | Data source | Years | Units | Resolution | Location | Rehab | New | Food |
|---|---|---|---|---|---|---|---|---|---|
| Maximum temperature | tempMax | AGCD | 1996-2021 | ºC | 5 km | ✓ | ✓ | ✓ | |
| Minimum temperature | tempMin | AGCD | 1996-2021 | ºC | 5 km | ✓ | ✓ | ✓ | |
| Difference in temperature (max – min) | tempDiff | AGCD | 1996-2021 | ºC | 5 km | ✓ | ✓ | ✓ | |
| Total precipitation | prec | AGCD | 1996-2021 | mm | 5 km | ✓ | ✓ | ✓ | |
| Precipitation anomaly | prec_anom | AGCD | 1996-2021 | - | 5 km | ✓ | ✓ | ✓ | |
| NDVI | ndvi | AGCD | 1996-2019 | - | 5 km | ✓ | ✓ | ✓ | |
| Soil moisture at root level (1 m) | soilMoisture | AGCD | 2000-2021 | % | 5 km | ✓ | ✓ | ✓ | |
| Vapor pressure at 9:00 AM | | AGCD | 1996-2021 | hPa | 5 km | ✓ | ✓ | ✓ | |
| Solar exposure | solarExposure | AGCD | 1996-2021 | $MJ/m^2$ | 5 km | ✓ | ✓ | ✓ | |
| Potential evapotranspiration | evapPotential | AGCD | 2000-2021 | mm | 5 km | ✓ | ✓ | ✓ | |
| ONI | ONI | NOAA | 1996-2021 | - | - | ✓ | | ✓ | ✓ |
| SOI | SOI | NOAA | 1996-2021 | - | - | ✓ | | ✓ | ✓ |
| SAM | SAM | BAS | 1996-2021 | - | - | ✓ | | ✓ | ✓ |
| Forest cover | perc_forest | DLCD | 2001-2015 | % | 250 m | ✓ | ✓ | ✓ | |
| Urban area | perc_urban | DLCD | 2001-2015 | % | 250 m | ✓ | ✓ | ✓ | |

| | | | | | | | | |
|---|---|---|---|---|---|---|---|---|
| Crop area | perc_crop | DLCD | 2001-2015 | % | 250 m | ✓ | ✓ | ✓ |
| Pasture area | perc_pasture | DLCD | 2001-2015 | % | 250 m | ✓ | ✓ | ✓ |

**Table S2. Details of lagged variables used as environmental features.** We calculated lagged versions of several variables shown in Table S2. For each variable, we show the variable name, the units each lagged variable is in, the code typically associated with the variable if it is different from the variable name, the particular months that each variable was lagged, and the method used to create the lagged variable (SD - standard deviation, Sum - sum of values, Value - value from that particular index *n* months prior).

| **Name** | **Units** | **Variable code** | **Lags (in months)** | **Lag method** |
|---|---|---|---|---|
| Maximum temperature | - | tempMax_[lag] | 2, 3, 6, 9, 12, 15, 18, 21, 24 | SD |
| Minimum temperature | - | tempMin_[lag] | 2, 3, 6, 9, 12, 15, 18, 21, 24 | SD |
| Difference in temperature (max – min) | - | tempDiff_[lag] | 2, 3, 6, 9, 12, 15, 18, 21, 24 | SD |
| Total precipitation | mm | prec_[lag] | 2, 3, 6, 9, 12, 15, 18, 21, 24 | Sum |
| Precipitation anomaly | - | prec_[lag]anom | 2, 3, 6, 9, 12, 15, 18, 21, 24 | SD |
| NDVI | - | ndvi_[lag] | 2, 3, 4 | SD |
| Solar exposure | $MJ/m^2$ | solarExposure_[lag] | 2, 3, 6, 9, 12, 15, 18, 21, 24 | Sum |
| ONI | - | ONI_[lag] | 2, 3, 6, 9, 12, 15, 18, 21, 24 | Value |
| SOI | - | SOI_[lag] | 2, 3, 6, 9, 12, 15, 18, 21, 24 | Value |
| SAM | - | SAM_[lag] | 2, 3, 6, 9, 12, 15, 18, 21, 24 | Value |

#### [B.ii] **Component model labels**

We constructed the multiple component models within our multi-scale modeling framework with data of varying spatial resolution (Table S2) and spatial extents (Fig. S1). The variation in resolution and extent was driven by different questions being answered by different component model (e.g., black flying fox distributions cover a larger area than spillovers have been reported) and by data limitations.

Data derived from a combination of previously published datasets, publically available datasets and data made available for this study. The roost distribution model was constructed from multiple datasets. The National Flying Fox Monitoring Program (NFFMP) recorded data from Queensland (public; National Flying Fox Monitoring Program, 2020) and New South Wales (shared by CSIRO Environment). The dataset of active roost sets (ARS) was used in the roost distribution model and new overwintering roosts (Eby et al. 2022a). The Wildlife Information, Rescue and Education Service (WIRES) Mid North Coast and Northern Rivers generously made summary data available, this is also publicly available on the associated FigShare (https://figshare.com/s/ddb5a1584609b20f6596). Lastly, the dataset of nectar shortages (aka NECT) was used to model acute food shortages (Eby et al. 2022b).

**Table S3. A summary of the component model data and their characteristics.** Along with the model name, we show the number of rows in each dataset, the minimum and maximum year encompassed in each model, the temporal and spatial scale of the environmental data used, and what source these data points were sourced from.

| Model | Data points (#) | Temporal extent (yrs) | Temporal scale | Spatial resolution | Data sources |
|---|---|---|---|---|---|
| Roost Distribution Model | 18,861 | 1996 - 2021 | month | 5 km | NFFMP, ARS |
| Roost count model | 53 | 2008 - 2020 | quarter | Study area | NFFMP |
| Bat Rehab | 10,150 | 2005 - 2020 | month | Town polygons | WIRES |
| New overwinter roosts | 195 | 2002 - 2019 | month | 5 km | ARS |
| Food Shortage | 267 | 1998- 2020 | month | One area | NECT |

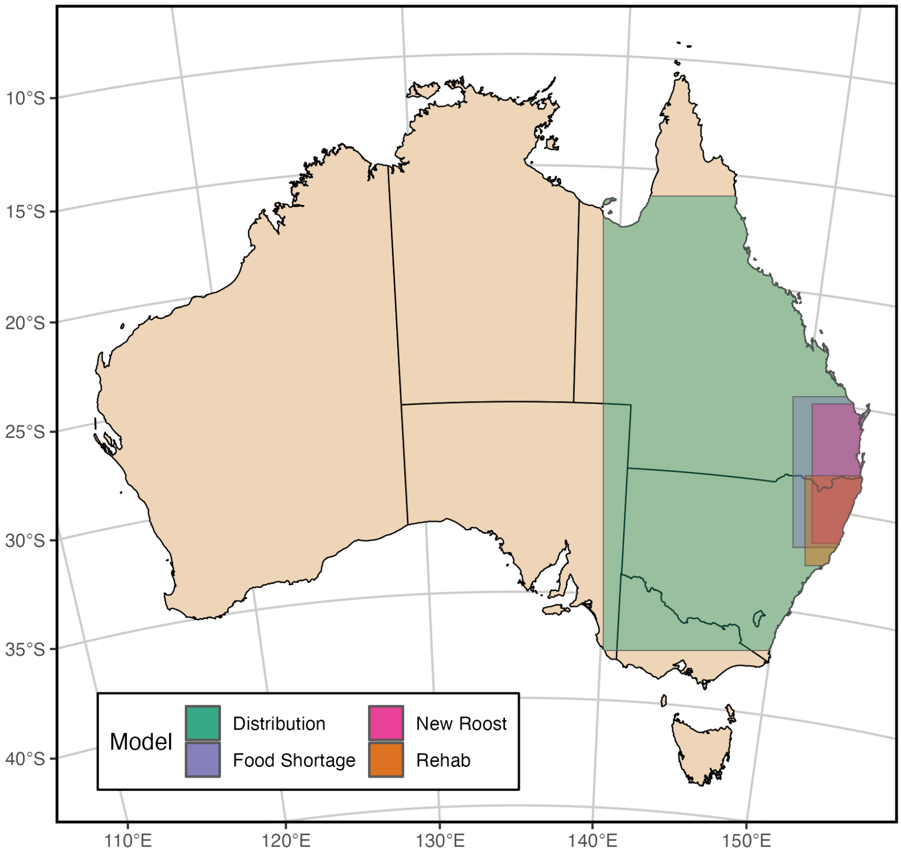

**Figure S1. Spatial extent of each component model used in our multi-scale framework.** The rectangles represent the bounding boxes that contain all points used in training each of these component models.

### [C] COMPONENT MODEL DETAILS

Our multi-scale model integrates the predictions from three different component stress models and also takes advantage of modeling performed to predict the suitability of roosts across the landscape and number of roosts occupied at a particular point in time (Table S3).

For each component model except for the estimation of the number of occupied roosts, we conducted parameterization and evaluation in the same fashion. All modeling was done using the gbm (version 2.1.8.1) and caret (version 6.0-86) packages in R (Kuhn 2020, Greenwell et al. 2022, R Core Team 2024), with the exception of the food shortage model that used XGBoost (Chen et al. 2023). Model parameterization was done using a grid search approach that iteratively searched through combinations of parameter values for learning rate (eta), max depth (depth of interactions), and minimum number of observations in a node. We used 80% of the data to train a model using a particular combination of these parameter values and 20% of the data to test the model. For choosing the final set of parameters to evaluate and train each component model, we used test area under the curve (AUC), test root-mean-square error (RMSE), and a visual comparison of deviance curves. For the models discussed below, we present the final parameters chosen and the number of trees run for each model.

We evaluated models using a bootstrap approach with 50 bootstrap iterations to determine average test AUC, calculate relative importance for each variable used, and create partial dependence curves. It is possible that patterns in the environmental data used here can affect the accuracy of our models, irrespective of their relationship with our labels. To investigate this, we used a null model approach by shuffling the target labels for the dataset and running the model for 50 bootstrap iterations using the same parameters. Using the test AUC from these null model iterations, we corrected our test AUC by subtracting the amount above 0.5 (the value expected for a model performing equal to random) derived from the null model approach. For each bootstrap iteration, we trained the model on 80% of the data and tested using 20%. For each model discussed below, we report the average corrected test AUC and show figures representing the variable relative importance and partial dependence results.

#### [C.i] *Roost distribution*

*Data*

We collated data on quarterly roost surveys from the National Flying Fox Monitoring Program (NFFMP) in Queensland and New South Wales. Roost survey dates are exact and were summarized for the month in which they occurred. NFFMP surveys of flying fox roosts have been conducted quarterly, recording the approximate population size and species identity during the daytime in coordinated observation windows. These roost data were supplemented by another set of roost surveys (Eby et al. 2022a). This second dataset did not have monthly observations, so we selected only roosts known to have overwintering bats, making the assumption that bats at these roosts were present throughout the winter. These roost data were replicated to cover all winter months (June, July, and August) and added to the NFFMP data to give a total of 18,861 roost records (Fig. S3). We defined presence data as roosts that were observed with at least one black flying fox (*Pteropus alecto*), whereas absences were roosts observed in surveys, but not containing black flying foxes (i.e., either being observed as occupied only by other species, or not occupied at all). Overall, there were 10,622 roost-timepoints occupied by *P. alecto* and 8,239 roost-timepoints with no *P. alecto* individuals observed from 1906-2021 (Table S3). We used the terra (version 1.5-21) and sf (version 1.0-7) packages in R version 4.1.0 to extract environmental data using point coordinate information associated with these roosts (Pebesma and Others 2018, Hijmans 2022, R Core Team 2024). For land cover classifications, we were interested in the percentage of each land cover class within the typical foraging area. We buffered coordinates by 20 km before extracting land cover data within this buffered region from the Dynamic Land Cover Dataset of Australia (Lymburner et al. 2015) and summarizing each land cover class (forest, urban, crop, pasture) as a percentage. Future models could test accuracy of these assumptions and improve model predictions by using scalable spatial scales and analyzing the most appropriate spatial buffers for different settings (i.e. Brock et al. 2019)

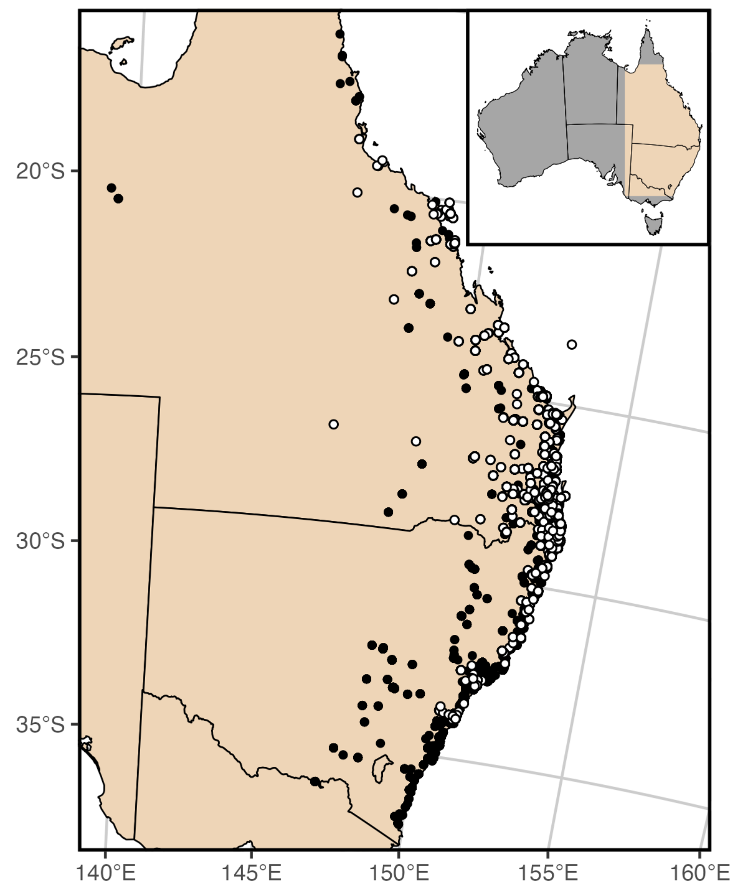

**Figure S2.** Occurrence data used in the roost distribution model from the National Flying Fox Monitoring Program and Eby et al. 2023. White circles represent roosts where a black flying fox has been surveyed at a point in time while black circles represent roosts with no black flying foxes found at the time of survey. The inset map in the top right shows the study range (beige) in the context of the rest of Australia.

*Model construction and hyperparameters*

We fit a classification model using generalized boosted regression via the gbm package (version 2.1.8.1) in R (Greenwell et al. 2022). Parameterization via grid search determined the most appropriate combination of hyperparameters to be an eta of 0.01, max depth of 4, and number of minimum observations in a node to be 2 with 200,000 trees. The 'best fit model' described below refers to the model fit with these hyperparameter set.

*Best fit model*

Model evaluation was performed using 30 iterations of five-fold cross validation, calculating AUC, relative importance scores, and metrics used to construct partial dependence plots from these bootstrap iterations. We compared these results to 30 iterations using randomly shuffled labels. We used the results from this null distribution of AUC values to correct for any inflation in evaluation caused by underlying patterns in the predictor data. This mean corrected AUC across 30 iterations of bootstrapping was 0.867. The confusion matrix showed that 89.5% of records were classified correctly using a threshold of 0.5.

The model of black flying fox roost occupation fit using our best performing set of hyperparameters as determined by our grid search approach was predicted by several temporal and spatial environmental variables. Full details of the partial dependence of the top 25 (out of 91) most important variables are in Figs. S3 and S4. The top four variables were all percentage of land cover (including urban, pasture, crop land, and forest) within a 20km radius foraging area surrounding the roost. Black flying fox roosts are more likely to be occupied in areas with higher urban cover, intermediate pasture and crop, and extremes of forest cover (high and low). The next most important variables included temperatures (standard deviation of maximum, minimum and range), cumulative solar exposure, and soil moisture (Fig. S3). Black flying foxes were more likely to occupy roosts that had low variability in temperature and higher soil moisture (Fig. S4). Interestingly, NDVI (greenness of vegetation) had variable impacts on roost occupation.

**Figure S3**. **Variable importance plot for the best fit roost distribution model. Variables are** arranged in descending order of importance. Relative importance is shown on the x-axis and variable name is shown on the y-axis. Points represent the mean importance value and bars represent the 95% confidence interval.

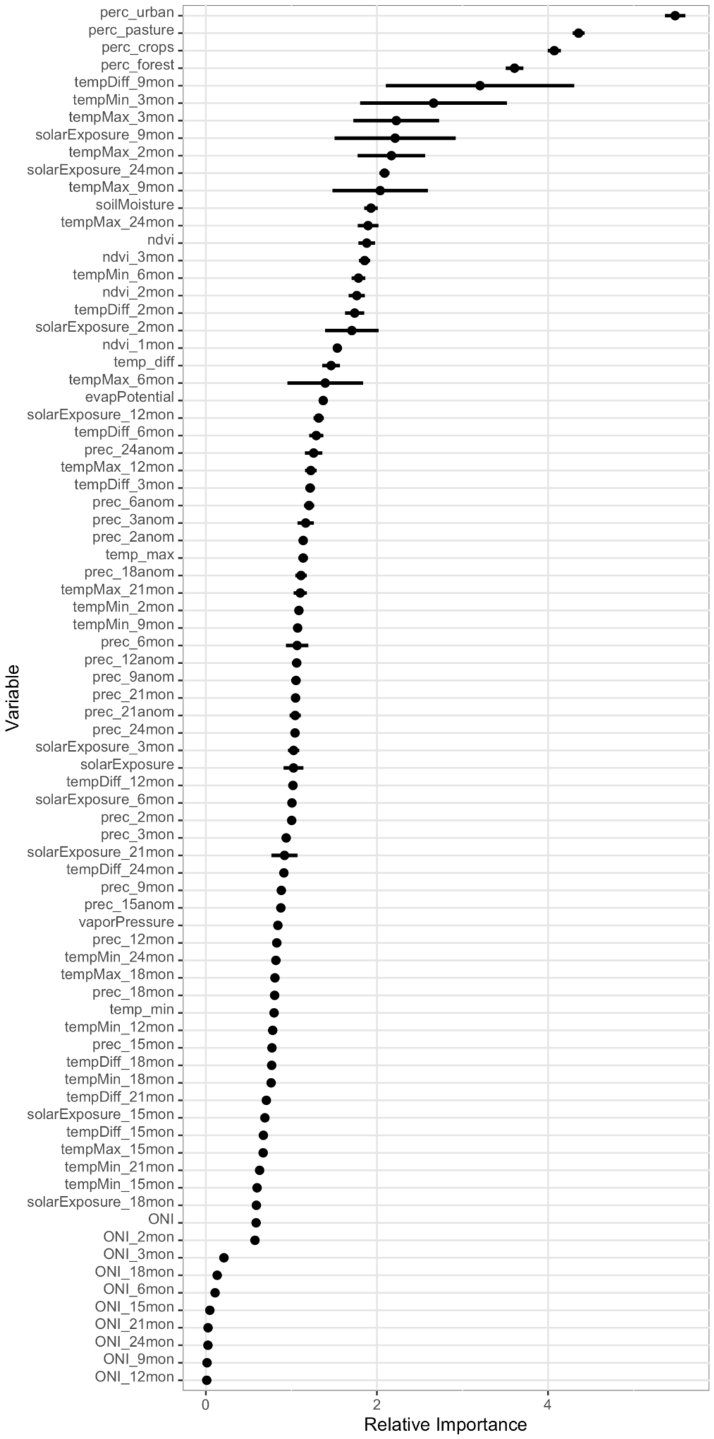

**Figure S4.** Partial dependence plots for the roost distribution model subset to the top 25 most important variables. For each panel, the left y-axis shows the marginal effect on prediction. The right y-axis shows the frequency of the variable value as represented by the histogram in the panel, and the x-axis represents the variable value. The bolded line in each panel represents the mean relationship of the variable's value to prediction while the shaded region is the 95% confidence interval from 30 bootstrap iterations.

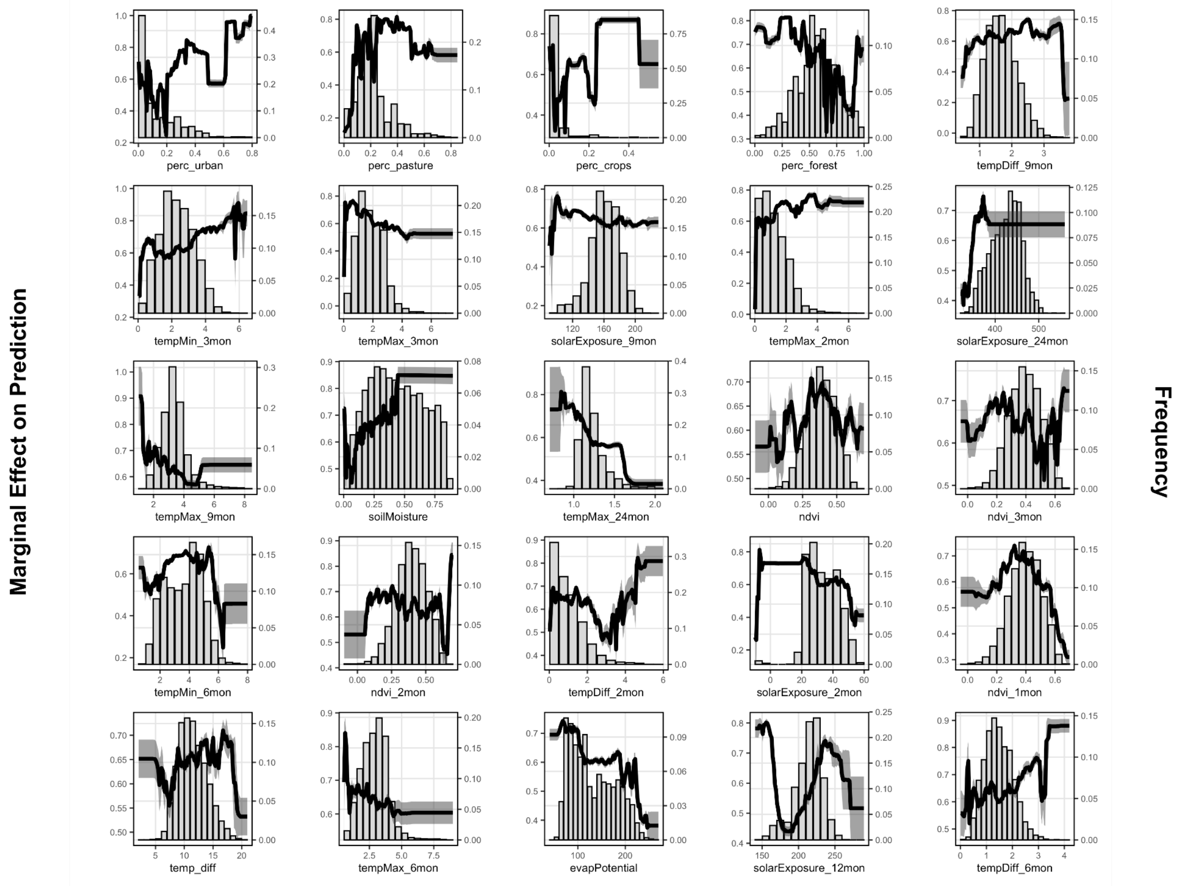

#### [C.ii] *Flying fox rehabilitation admissions*

We constructed a component model to reflect admission of flying foxes to rehabilitation facilities. This model was constructed to estimate probabilities of a bat being delivered to a rehabilitation facility, using summary intake data from two wildlife rescue branches.

*Data*

Data regarding a series of bats encountered by two Wildlife Information Rescue and Education Service (WIRES) branches between 2005 - 2020 were kindly provided by WIRES Northern Rivers and WIRES Mid North Coast WIRES (Table S3; dataset made public here: https://figshare.com/s/ddb5a1584609b20f6596). Using provided information about the specific locality and postcode each flying fox was collected in, we associated each record with polygons representing the boundaries of suburbs (in cities and larger towns) and localities (outside cities and larger towns) created by Geoscape Australia available from the Australian government via data.gov.au (https://data.gov.au/data/dataset/geoscape-administrative-boundaries). Some bat records were associated with public lands, so boundaries of these areas were obtained from the New South Wales National Parks and Wildlife Service Estate Database (https://datasets.seed.nsw.gov.au/dataset/nsw-national-parks-and-wildlife-service-npws-estate3f9e7) and the legal State Forest Boundaries of New South Wales (https://data.gov.au/dataset/ds-nsw-5b803c70-08c3-4bac-8f64-a0261dcbca12/details?q=). These data totaled to 5,075 bats recorded between 2005 and 2020 (Table S3).

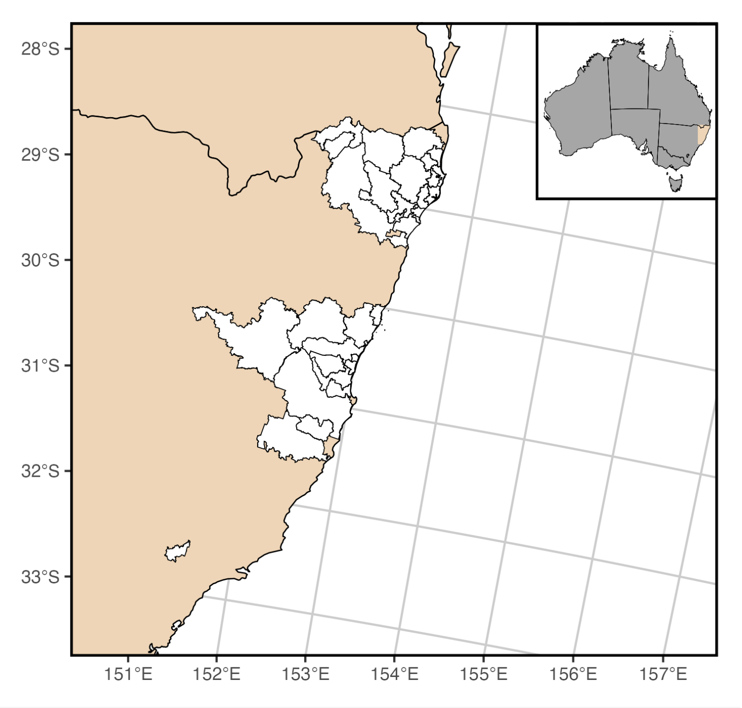

**Figure S5.** Polygons (white shapes) show the postcodes that bats delivered to rehabilitation centers were associated with. Presence and background data were derived from suburb and locality boundaries within these postcodes. The inset map in the top right shows the extent of these data (beige) in the context of Australia.

We determined an equal amount of random background data points by randomly choosing polygons within the postcodes associated with the rehabilitated bats. Preliminary results from initial models suggested overfitting to land cover data, so we filtered potential background points on the distribution observed in presence data to exclude postcodes composed mainly of pasture (>76.8% land cover) from selection. We also weighted background point selection to match the distribution of urban land cover seen within our rehabilitation data. Briefly, we determined the distribution of rehabilitated bat points in deciles of possible urban land cover (e.g., number of bats found in 0-10% urban land cover, number in 10-20%, etc.) and biased background point selection to match this distribution. Years were assigned to match the same distribution of years found in our bat records and months were assigned randomly. For all records (rehab and background data), we extracted environmental data using the specific locality polygons and took the mean value found within the polygon. All calculations of differences, anomalies, and lags were done using all extracted values before averaging.

*Model construction and hyperparameters*

We fit a classification model using generalized boosted regression with the gbm package in R. Parameterization using grid search and careful examination of evaluation statistics and deviance curves determined the best parameter combination to be an ETA of 0.0001, max depth of 2, number of minimum observations per node of 10, and number of trees of 150,000.

*Best fit model*

Model evaluation was performed similarly to the roost distribution model using 50 bootstrapped iterations and correcting the AUC using the same number of iterations run with shuffled labels. The mean corrected AUC for the bat rehabilitation model was 0.876.

The best fit model of flying foxes admitted to rehabilitation was predicted by spatial characteristics and lagged temporal environmental features (Figs. S6 and S7). The top three variables were all percentage of land cover (including urban, pasture, and forest). Flying fox rehabilitation admissions were more common in postcodes with extremes of urban cover (very low and high), and unlikely in areas with low pasture and forest cover. The next most important variables included the standard deviation of maximum temperature and cumulative precipitation over the previous 24 months. Rehabilitation admissions were more common when the previous 24 months had lower variation in maximum temperatures and were on average wetter. The next several variables were environmental features varying from contemporaneous conditions to 24 month lagged variables.

**Figure S6.** Plot of variable importance for the bat rehabilitation model organized in descending order of importance. All variables with relative importance greater than 0.75 are shown. Relative importance is shown on the x-axis and variable name is shown on the y-axis. Points represent the mean importance value and bars represent the 95% confidence interval.

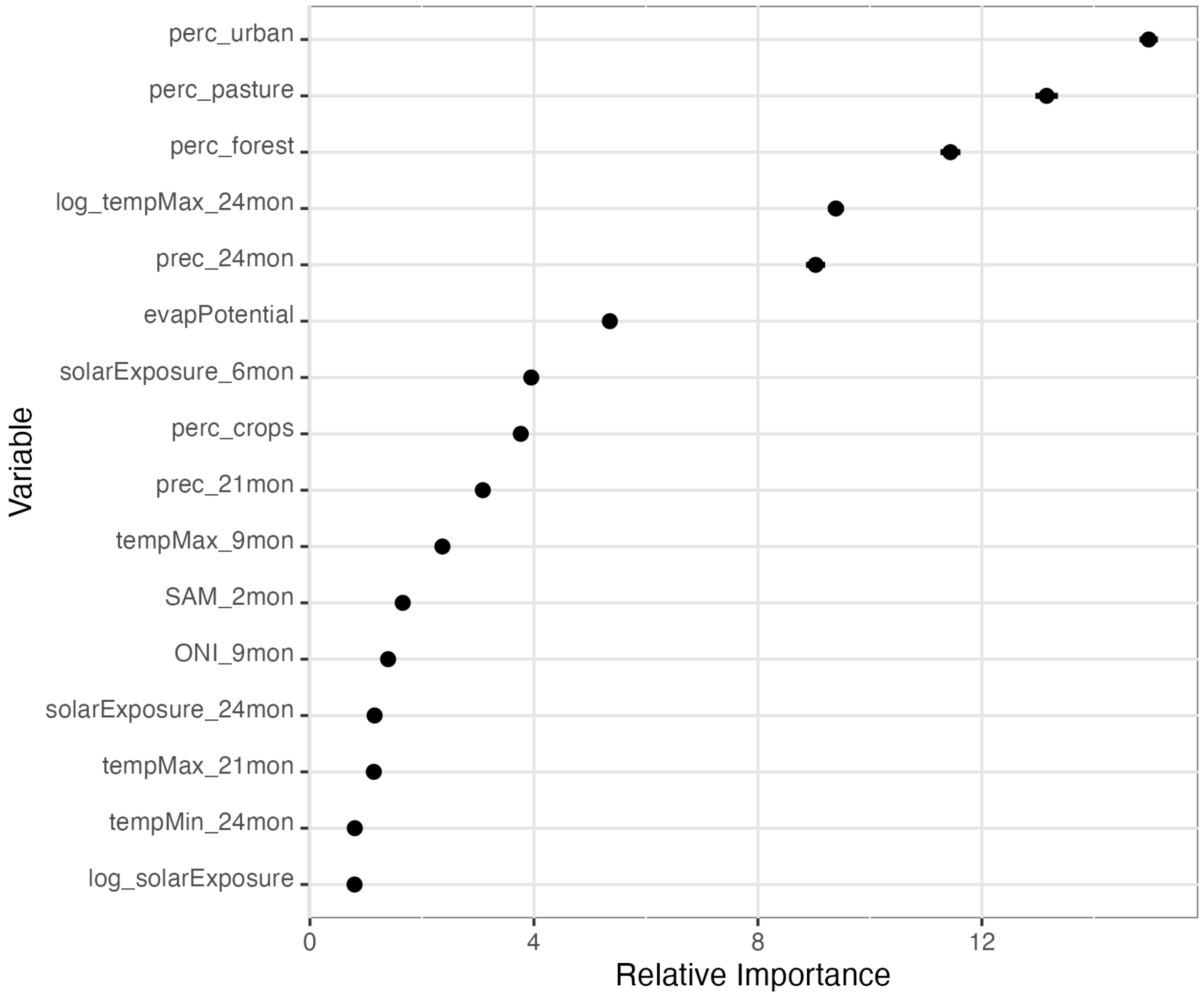
perc_urban
perc_pasture
perc_forest
log_tempMax_24mon
prec_24mon
evapPotential
solarExposure_6mon
perc_crops
prec_21mon
tempMax_9mon
SAM_2mon
ONI_9mon
solarExposure_24mon
tempMax_21mon
tempMin_24mon
log_solarExposure
Variable
0
4
8
12
Relative Importance

**Figure S7.** Partial dependence plots for the bat rehabilitation model subset to all variables with a relative importance greater than one. For each panel, the left y-axis shows the marginal effect on prediction, the right y-axis shows the frequency of the variable value as represented by the histogram in the panel, and the x-axis represents the variable value. The bolded line represents the mean pattern while the shaded region is the 95% confidence interval from 50 bootstrap iterations.

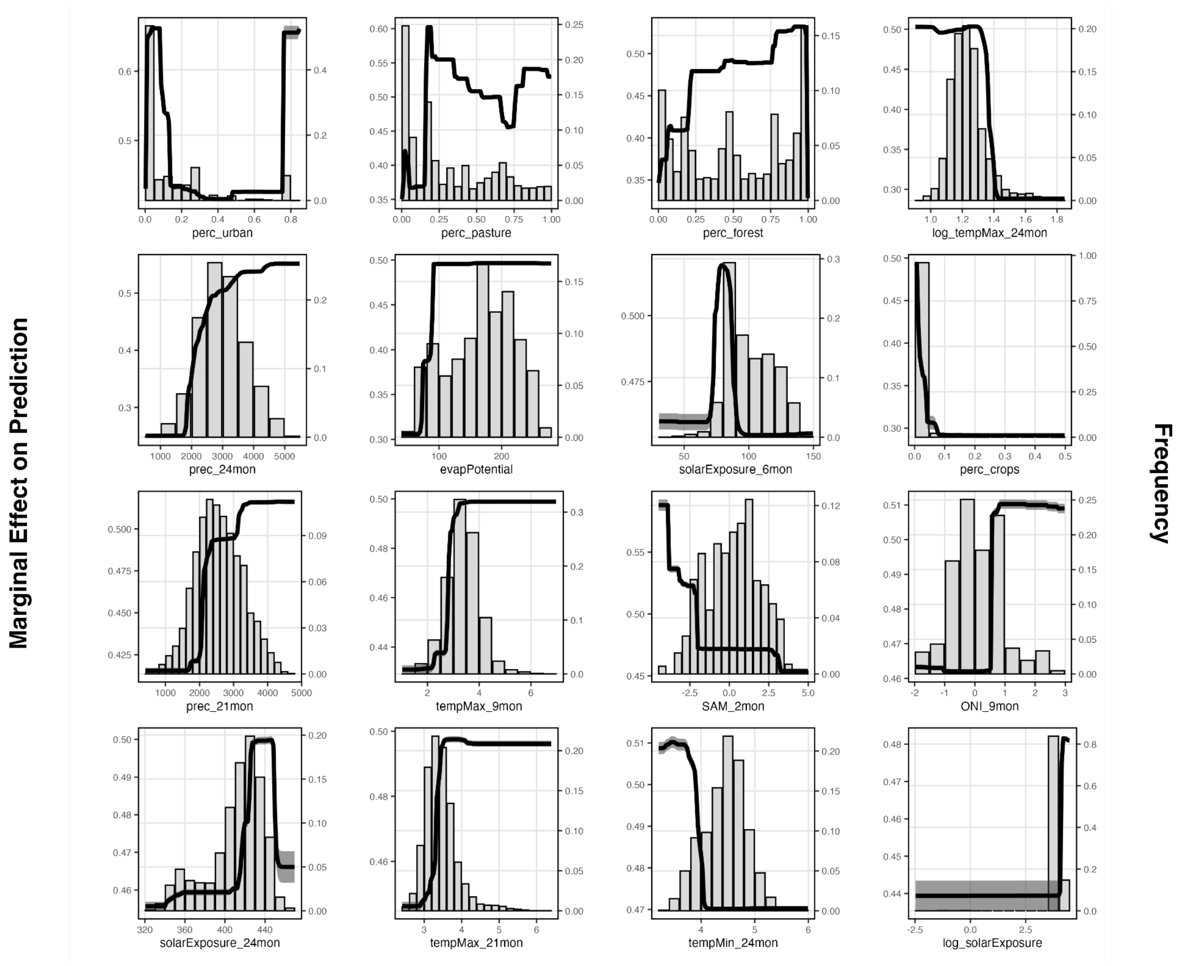

#### [C.iii] *Formation of new overwintering roosts*

We constructed a second component model to predict the probability of a roost in a particular location representing a new, fissioned roost. New overwintering roosts are hypothesized to be a stress response (Becker et al. 2023, Eby et al. 2023) and as such these locations may be more likely to have bats experiencing physiological stress.

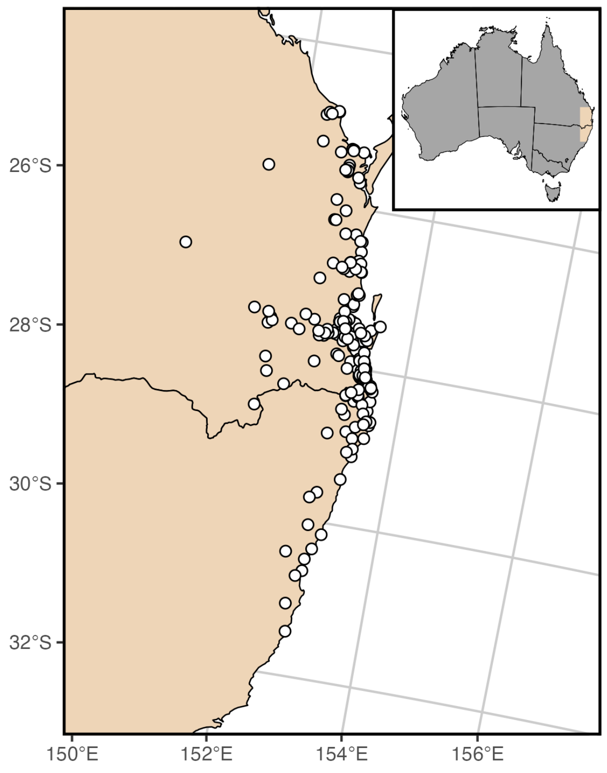

**Figure S8.** Occurrence points used in training our model to detect new roosts depicted as white filled circles. The inset map shows the study extent for this model (beige) in the context of Australia.

Using active roost survey data with years of formation documented previously (Eby et al. 2022a), we restricted the dataset to roosts that formed between 2002 – 2019 (Fig. S8). These newly formed roosts can represent splits in larger roosts or the movement of bats in search of new food sources (Tait et al. 2014, Páez et al. 2018). As the exact date of formation of a new roost is uncertain, we evaluated two different timepoints in years with acute food shortages: 1) the midpoint of an associated acute food shortage event or 2) last month of an associated acute food shortage event. In years without acute food shortage, we assume the roost was formed in August (last month of winter). We also compared several different methods of selecting background data after initial results (using non newly formed roosts as our contrast class) showed a severe urban bias towards classifying new roosts and suggested overfitting. To remedy this, we used the geographic extent of the surveyed roosts to constrain selection, we further removed waterbodies and areas within 20 km of a surveyed roost from consideration as a selected background point. We created three different sets of background points: those randomly chosen within this area, randomly chosen points in this area that matched the pattern of percent coverage by urban area found in the newly formed roosts, and a selection of points in the area guided by the results of our roost distribution model. To deal with a paucity of available points, the distance from surveyed roost buffer was reduced to 1 km during the selection of urban biased and roost distribution guided background points. For the background points selected by using the roost distribution model results, we calculated the true skill statistic (TSS) using newly formed roosts and urban biased background points. The suitability value corresponding to the maximum TSS was chosen to threshold roost suitability predictions and guide the selection of random points within our study area. Each of the points was randomly assigned a year from the distribution of years contained in the full dataset and all points were assigned the month of August. After a comparison of model results using each of these sets of background points, we ultimately chose to use background points selected using roost distribution suitability values. Additionally, we chose to set the timing of new roost formation to the last month of an associated food shortage as this performed better at classifying new roost formation from the background. These decisions minimized overfitting while improving accuracy of the model as measured by AUC and sensitivity as measured using a confusion matrix.

*Model construction and hyperparameters*

All comparisons were done using classification with generalized boosted regression with the gbm package in R. Depending on the background points used, model corrected AUC ranged from 0.648 (comparison with non newly formed roosts) to 0.954 (comparison with random background points).

*Best fit model*

The mean corrected AUC of the model using background points selected by roost distribution suitability values was 0.9 with final parameters of an ETA of 0.0001, maximum depth of 2, number of minimum observations in a node of 2, and best number of trees of 123,533. A confusion matrix showed that 98.462% of new roosts were correctly classified (192/195).

The best fit model of new roost formation was predicted by several temporal and spatial environmental variables. The top two variables (percentage urban land cover and standard deviation of maximum temperature in the previous two months) had high relative importance compared to the rest of the variables. New roosts are more likely in higher urban cover and when variation in maximum temperature is lower over the preceding two months. They are also more likely to form when the Southern Annular Mode (SAM) is low 15 months prior. Several other environmental features - such as soil moisture, precipitation and evaporation potential are relatively important in predicting new roosts. The most important variables and their marginal effects on prediction are shown in Figures S9 and S10.

**Figure S9.** Plot of variable importance for the new roost model organized in descending order of importance. Relative importance is shown on the x-axis and variable name is shown on the y-axis. Points represent the mean importance value and bars represent the 95% confidence interval.

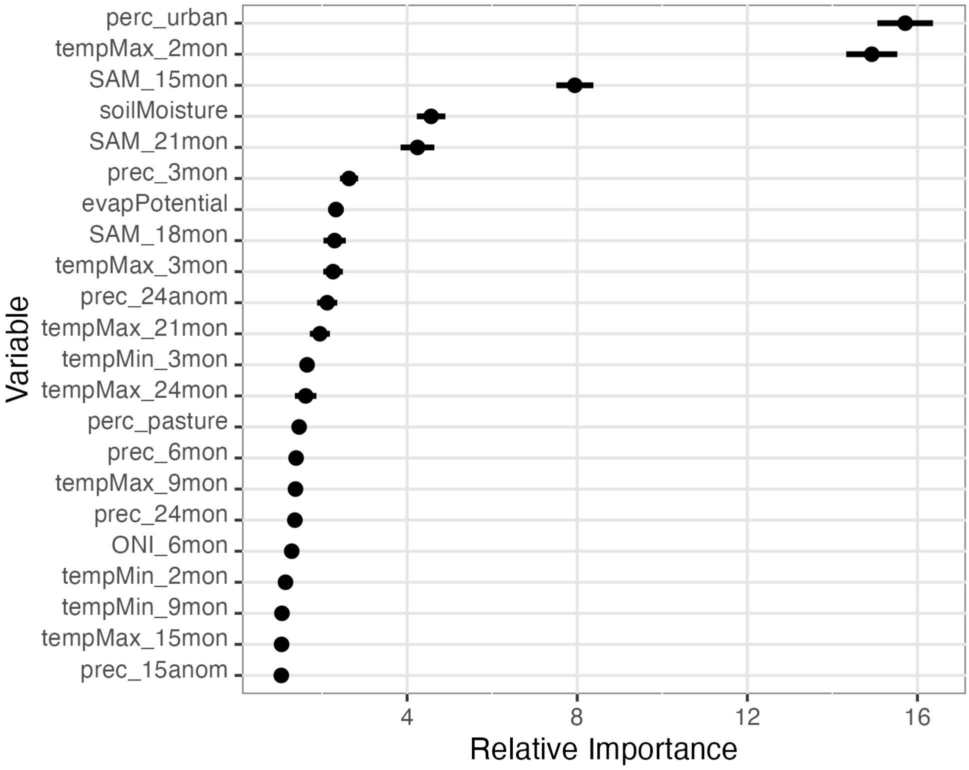

**Figure S10.** Partial dependence plots for the new roost model subset to all variables with a relative importance greater than one. For each panel, the left y-axis shows the marginal effect on prediction, the right y-axis shows the frequency of the variable value as represented by the histogram in the panel, and the x-axis represents the variable value. The bolded line represents the mean pattern while the shaded region is the 95% confidence interval from 50 bootstrap iterations.

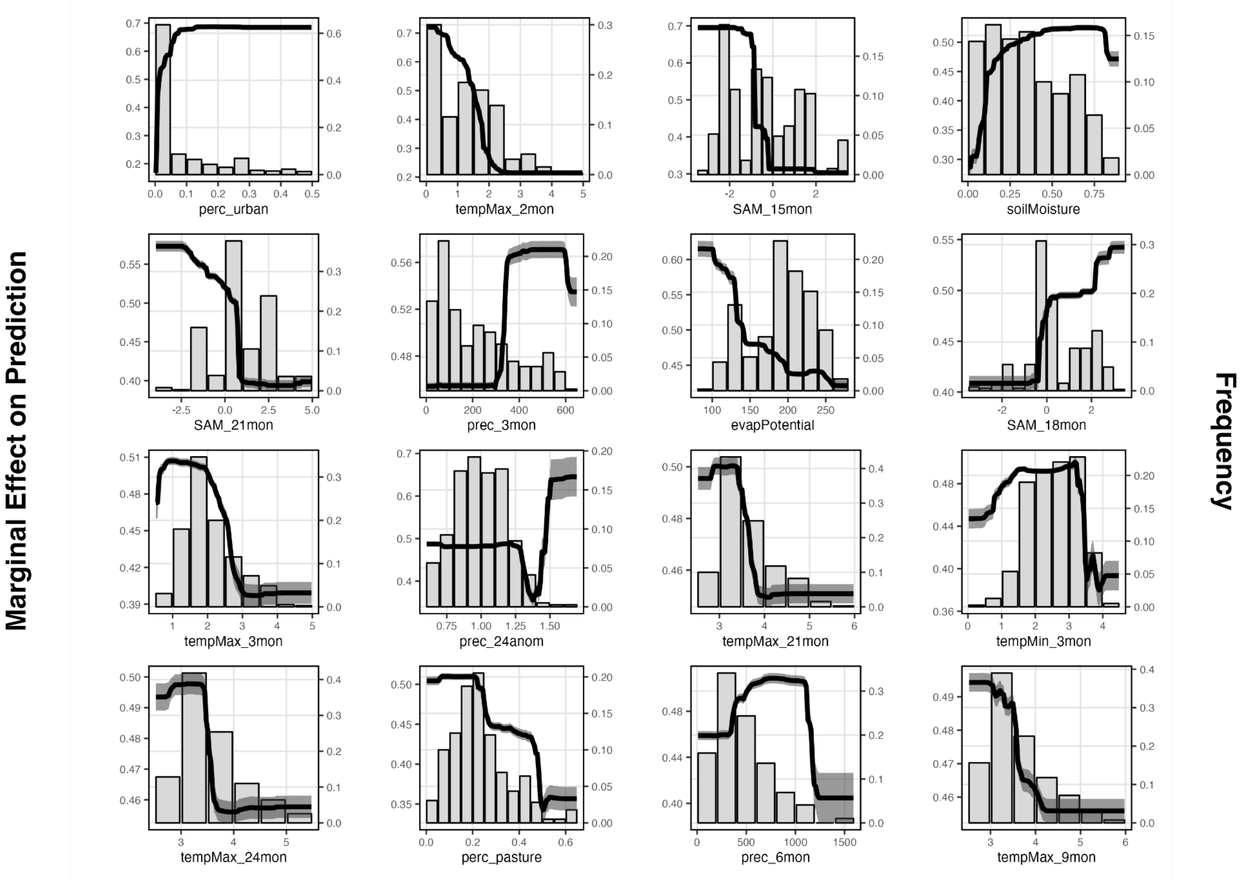

#### [C.iv] *Acute food shortage*

We constructed a third component model to represent host condition. This model was constructed to estimate probabilities of an acute food shortage, modeled using regional nectar shortage data.

*Data*

To model acute food shortage, we used records of nectar shortage from Eby et al. (2023) that range from 1998 until March 2020 (Table S3, Eby et al. 2022a). Based on surveys of apiarists, the authors classified their study area in the subtropics of eastern Australia (including southeast Queensland and northeast New South Wales) as having a nectar shortage or not. Following the analysis in Eby et al. (2023), we attached data summarizing ONI, SOI, and SAM at the time of the survey and at various time periods up to two years before the time of the survey (Table S1).

*Model construction and hyperparameters*

We used XGBoost to create a classification model of months of food shortage using the xgboost package in R (Chen et al. 2023). We parameterized the model using grid search, evaluating model combinations using logloss. The model parameter combination chosen was an ETA of 0.001, max interaction depth of 4, subsampling ratio of 1, column ratio to be sampled by tree of 0.5, minimum child weight of 1, gamma of 0.5, and number of trees of 15,000. We evaluated the model using 100 iterations of four-fold cross validation.

*Best fit model*

The corrected mean AUC after 100 iterations of bootstrapping was 0.811. The best fit model of food shortage was predicted by ONI, SAM, and SOI over different timescales. Three variables had high absolute mean SHAP values (a measure of relative contributions to model). These included ONI lagged by 9 months and 21 months and SAM lagged by 9 months. Interestingly, food shortage was more likely when there were higher ONI in the 9 months prior but lower ONI in 21 months prior. Food shortage was also more likely when SAM was higher 9 months prior. Several other lags and indices were included in the best fit model - but SOI and SAM over longer time scales had low SHAP values. Full details of the variable importance are in Figure S11 and partial dependence of the top 6 variables are shown in Figure S12.

**Figure S11.** A plot of variable importance for the food shortage XGBoost model with the mean absolute SHAP value on the x-axis and the variable name on the y-axis. The points represent the mean absolute SHAP values from 100 bootstrap iterations, while the line represents the extent of absolute SHAP values found from the same bootstrap iterations.

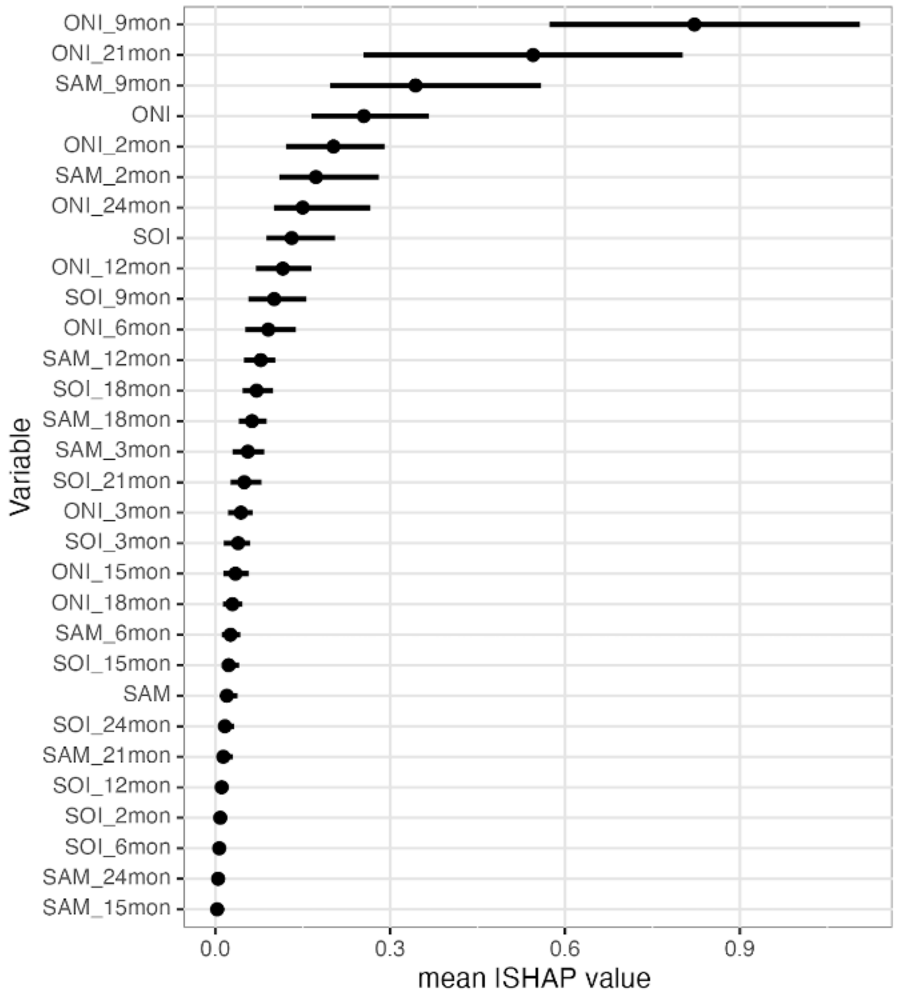

**Figure S12.** A subset of partial dependence plots from the top six most important variables as determined by mean absolute SHAP value. The thick blue line and thin gray lines represent the mean marginal effect of that variable on prediction and the pattern observed in each bootstrap iteration, respectively. For each panel, the x-axis shows the variable units and the y-axis represents the change in Shapley score. A rug plot is shown on the x-axis depicting the distribution of values found in the training dataset.

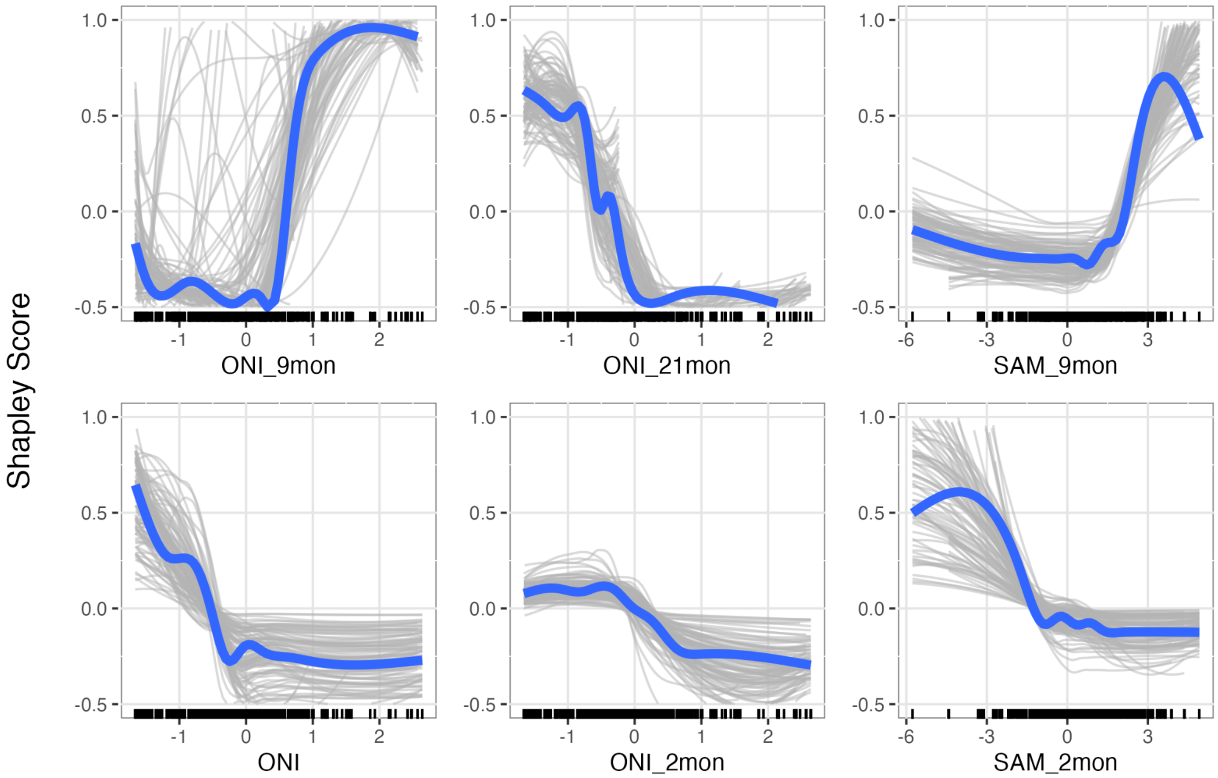

### [D] MULTI-SCALE MODEL CONSTRUCTION AND VALIDATION

#### [D.i] *Roost counts model*

For determining the number of roosts to estimate prevalence values at each monthly time step in our multi-scale model, we used a generalized additive model (GAM) to predict quarterly roost counts within our study area. We summarized the National Flying Fox Monitoring Program roost survey data to determine the number of roosts with black flying foxes found per quarter within the surveyed area, truncating these data to years that had all quarters surveyed (2008-2020). With these data, we used the *mgcv* package in R (Wood and Augustin 2002, Wood and Wood 2015) to fit a GAM to the quarterly roost count data using a cyclic cubic spline for the quarter and a cubic spline for the surveyed timepoint (53 unique timepoints surveyed). Model comparison indicated a model with uncorrelated errors was appropriate (following Simpson 2014).

#### [D.ii] *Model uncertainty*

We incorporated uncertainty in each of the component models by using 1,000 iterations of bootstrapping for each component. From this, each of the multi-scale model iterations used a separate bootstrapped component model. For each of these bootstrap iterations, we randomly sampled with replacement from the positive labels (i.e., roosts with black flying foxes, rehabilitation admissions of bats, newly formed roosts, and food shortage events) for the same number of presence points in each dataset. The predicted number of occupied roosts [D.i] were randomly distributed for a given month and weighted the probability of choosing a given grid cell using the roost occupancy model output. Each roost was assigned a unique foraging area by considering surrounding roosts and creating Voronoi tessellations to avoid overlapping foraging. The maximum foraging distance from a roost was 50 km - all tessellations were truncated to this distance if necessary. This divided areas between roosts when they were nearby while preventing unrealistically distant areas from being included.We also calculated a new round of background data for each iteration, taking care to apply any of the filters to reduce overfit that were used in evaluating the component models described above. We saved all component model objects from each iteration for use within our multi-scale modeling framework with all model objects found in a Figshare repository (https://figshare.com/s/ddb5a1584609b20f6596).

#### [D.ii] *Time lags*

When comparing predicted prevalence from the multiscale model and observed Hendra prevalence values from Field et al (2015), we found negative correlations (Table S4). Previous studies have demonstrated temporal lags between stress and subsequent spillovers in this system, potentially due to the long-time scales affecting eucalypt phenology (Hudson et al. 2011, Giles et al. 2016, Eby et al. 2023). We tested multiple combinations of this cumulative lag: no lag, 3, 6, 9, and 12 months lag. Our evaluation metrics comparing predicted prevalence from multi-scale models to observed prevalence in roosts (Field et al. 2015) showed that the 12 month cumulative lag performed the best for each of the model components (Fig. S14).

**Table S4.** Performance of single and multi-scale model combinations. In the Model column, 'All condition models' refers to the multi-scale model with rehabilitation, new roosts, and food shortage models. The null model only includes roost suitability without any condition modifiers. All correlations are shown with no lags - correlation refers to mean Spearman rank correlation coefficients.

| Model | Correlation | RMSE |
|---|---|---|
| All condition models | -0.161 | 0.407 |
| Rehab | -0.005 | 0.271 |
| New Roost | -0.200 | 0.254 |
| Food shortage | -0.031 | 0.094 |
| Rehab + New Roost | -0.143 | 0.398 |
| Rehab + Food shortage | -0.029 | 0.288 |
| Food Shortage + New Roost | -0.222 | 0.271 |
| Null | 0.214 | 0.293 |

**Figure S14.** A comparison of RMSE results for each model combination and multiple cumulative lag lengths for the model subcomponents (food shortage, new roost, rehab), showing that the cumulative 12 month lag for food shortage minimized RMSE. The x-axis shows each of the models run (non-12 month lagged models have their lag lengths shown) while the y-axis shows the RMSE. Each point represents the RMSE of a validation site, with the horizontal line representing the weighted mean based on the number of times a site was sampled.

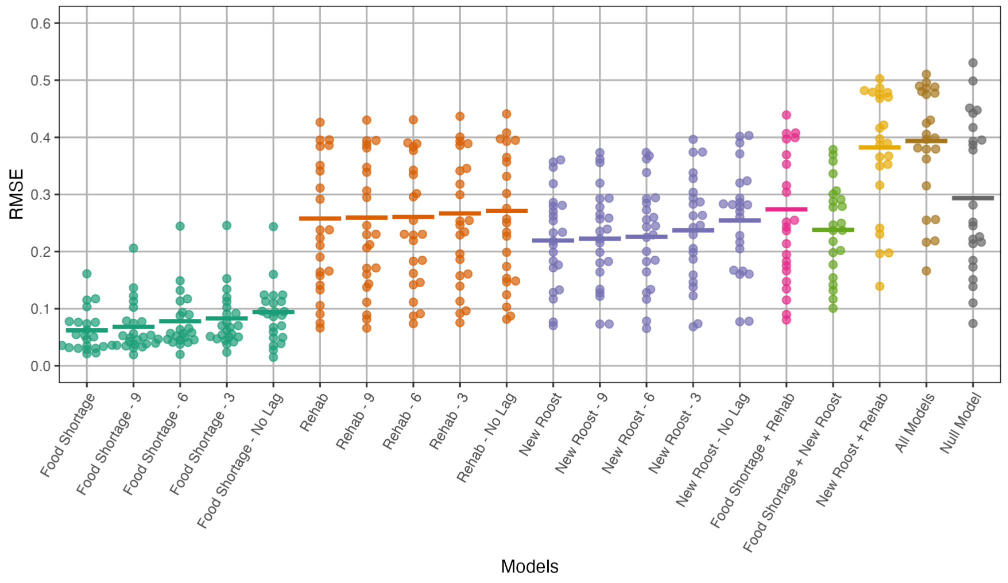

#### [D.iv] *Leave one out model description*

To better assess the influence of each of the models included in our multi-scale modeling process, we modeled predicted prevalence of HeV using all combinations of the new roost, rehabilitation, and acute food shortage models. This resulted in seven sets of predictions of monthly predicted prevalence from 2008 to 2020. Performance of each of these predictions was assessed using RMSE with observed data (Tables 1 and S3; Fig. S15). RMSE values were also normalized using the hydroGOF package in R using the difference between maximum and minimum values (Zambrano-Bigiarini 2020). This normalized RMSE (NRMSE) typically ranges from 0 to 100 unless major outliers exist. Spearman rank correlation was also calculated using the stats package in R (R Core Team 2021).

Predicted prevalence of HeV for each site was calculated by taking a 20 km buffer around the site coordinates. This takes into account a previous estimate for daily movement of *Pteropus alecto*. Within this buffered area, we used the predicted suitability of roosts to apply a threshold of 0.5 predicted suitability and remove estimated prevalence values associated with unsuitable habitat. For roost sites where all of the buffered area was considered unsuitable habitat based on this threshold, we kept areas matching the top 20% of suitability values within the buffered area and removed the rest. We also tried omitting sites with unsuitable habitat in a particular month from the analysis, but that only increased correlations slightly (+0.02).

The mean predicted prevalence value of the remaining habitat in a site's buffered area was kept as the predicted prevalence for that site for RMSE and correlation estimates. Variance in these predicted prevalences arises from the model iterations and is shown in Figure S16.

**Figure S15.** A comparison of RMSE results across each site for each of the multi-scale model combinations (depicted by color). Sites are arranged by latitude (with the most southern sites to the left) with the number of times the roost was sampled given in parentheses. The weighted mean of each model across all sites is given by the horizontal line.

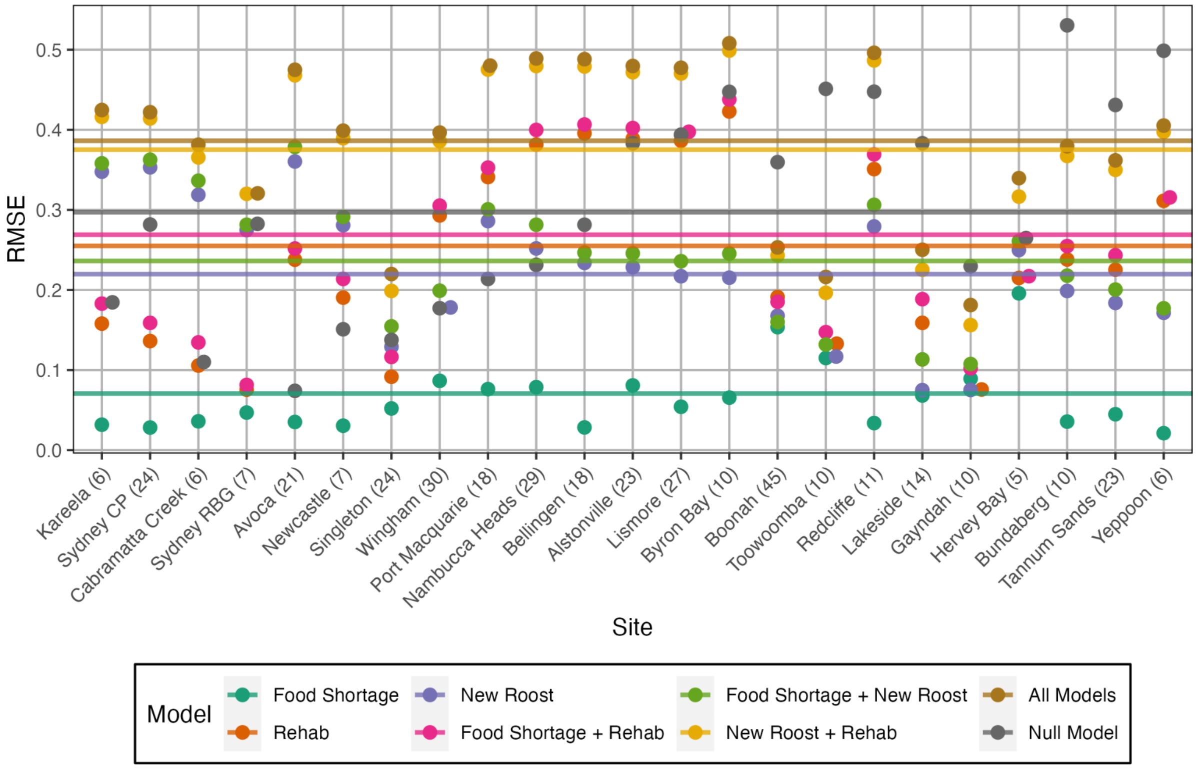

**Figure S16 (following page).** Monthly predicted HeV prevalence from 2011 until 2014 for our model using food shortage and the null model using locations for each of the Field et al. (2015) roosts. Sites are organized by latitude (with the most northern sites at the top). Light gray points represent estimates from the null model, while dark gray points represent predictions from the multi-scale model including acute food shortage. Larger, magenta points show observed HeV prevalence at each of the sites.

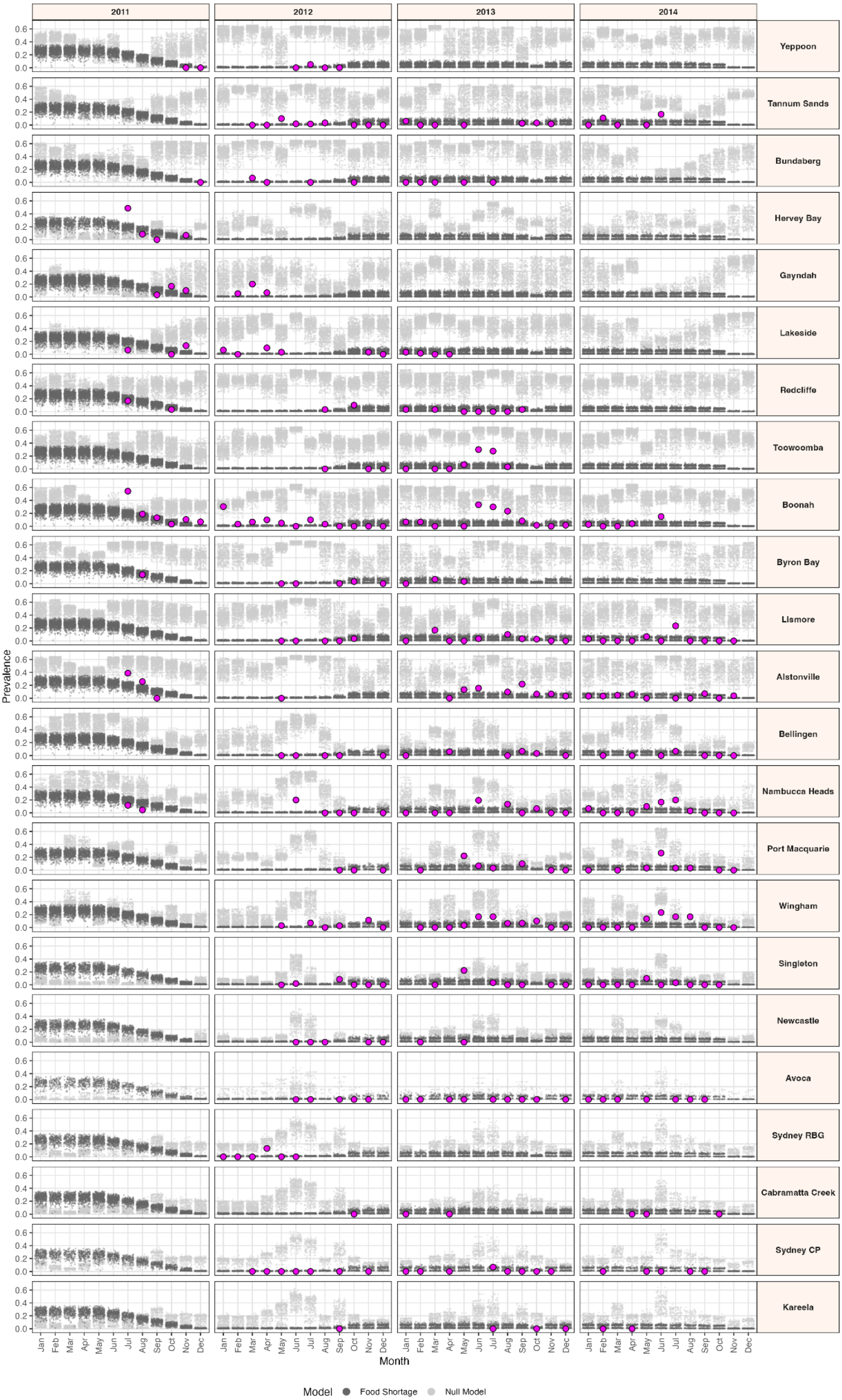

2011
2012
2013
2014
Yeppoon
Tannum Sands
Bundaberg
Hervey Bay
Gayndah
Lakeside
Redcliffe
Toowoomba
Boonah
Byron Bay
Lismore
Alstonville
Bellingen
Nambucca Heads
Port Macquarie
Wingham
Singleton
Newcastle
Avoca
Sydney RBG
Cabramatta Creek
Sydney CP
Kareela
Prevalence
0.0 0.2 0.4 0.6
Jan Feb Mar Apr May Jun Jul Aug Sep Oct Nov Dec
Month
Model
Food Shortage
Null Model
